# Expression of the *Bacillus thuringiensis vip3A* insecticidal toxin gene is activated at the onset of stationary phase by VipR, an autoregulated transcription factor

**DOI:** 10.1101/2022.04.23.489259

**Authors:** Haibo Chen, Emilie Verplaetse, Leyla Slamti, Didier Lereclus

## Abstract

The **V**egetative **i**nsecticidal **p**rotein Vip3A is produced by some *Bacillus thuringiensis* strains from the mid-log growth phase to sporulation. Although Vip3A is important for the entomopathogenicity of *B. thuringiensis*, the *vip3A* gene regulation is unknown. In the *B. thuringiensis kurstaki* HD1 strain, *vip3A* is carried by the pBMB299 plasmid, which is absent in the closely related strain *B. thuringiensis kurstaki* HD73. Using a transcriptional fusion between the *vip3A* promoter and *lacZ*, we observed that the HD73 strain is unable to express *vip3A*. This result suggests that a specific regulator is required for *vip3A* expression. Assuming that the regulator gene is located on the same plasmid as *vip3A*, we transferred the pBMB299 from the HD1 strain to the HD73 strain. We found that Vip3A was produced in the HD73 strain containing pBMB299, suggesting that the regulator gene is located on this plasmid. Using this heterologous host and promoter-*lacZ* transcription fusions, we confirmed that VipR is essential to activate *vip3A* expression at the onset of stationary phase. We demonstrated that *vipR* transcription is positively autoregulated and the determination of the *vipR* and *vip3A* promoters pinpointed a putative VipR target upstream from the Sigma A-specific −10 region of these two promoters. Surprisingly, this conserved sequence was also found upstream of *cry1I* and *cry2* genes. Finally, we showed that *vip3A* and *vipR* expression is drastically increased in a Δ*spo0A* mutant unable to initiate sporulation. In conclusion, we have characterized a novel regulator involved in the entomopathogenic potency of *B. thuringiensis* through a sporulation-independent pathway.

**Importance:** The insecticidal properties of *Bacillus thuringiensis* are mainly due to Cry toxins which form a crystalline inclusion during sporulation. However, other proteins participate in the pathogenicity of the bacterium, notably the Vip3A toxins that are produced from vegetative growth to sporulation. The VipR regulator that activates *vip3A* gene expression at the onset of stationary phase is positively autoregulated and analysis of the promoter region of the *vip3A* and *vipR* genes reveals the presence of a highly conserved DNA sequence. This possible VipR target sequence is also found upstream of the *cry2A* and *cry1I* genes, suggesting that Cry toxins can be produced before the bacteria enter sporulation. Such a result could allow us to better understand the role of Cry and Vip3A toxins during the *B. thuringiensis* infectious cycle in insects, in addition to the primary role of the Cry toxins in the toxemia caused by ingestion of crystals.

## Introduction

The *Bacillus thuringiensis* species includes a large number of Gram-positive sporeforming bacterial strains belonging to the *Bacillus cereus* group and distinguished by the production of parasporal crystal inclusions (1). Many strains of *B. thuringiensis* are entomopathogenic and their insecticidal properties are primarily due to the crystal proteins that consist of Cry and Cyt toxins, encoded by plasmid genes (2–4). The presence of *cry* genes is the common feature of all strains of the *B. thuringiensis* species, and the expression of most *cry* genes is dependent on the sporulation-specific factors Sigma E and Sigma K, which are functional in the mother cell compartment during sporulation. However, a few *cry* genes do not follow the same regulation pathway (5). They are also expressed in the mother cell or in a non-sporulating subpopulation during the sporulation process, but independently of the sporulation sigma factors. These are notably the *cry3* genes encoding toxins active against coleopteran insects (6) and the *cry* genes of strain LM1212 which are expressed under the control of a specific transcriptional activator, CpcR, encoded by a plasmid gene (7).

Different combinations of Cry toxins can be found in the crystal inclusions depending on the *B. thuringiensis* strains. As an example, the commercial strain *kurstaki* HD1 contains 6 *cry* genes: *cry1Aa, cry1Ab, cry1Ac, cry1Ia, cry2Aa* and *cry2Ab* (http://www.lifesci.sussex.ac.uk/home/Neil_Crickmore/Bt/). All these *cry* genes encode proteins active against lepidopteran larvae and confer a broad insecticidal spectrum within this insect order (8). However, the pathogenicity of *B. thuringiensis* towards some insects and other arthropods also depends on various chromosomal factors including those belonging to the PlcR virulence regulon (9, 10). Moreover, several *B. thuringiensis* strains harbor plasmid genes encoding insecticidal proteins other than the Cry toxins. It is notably the case of the Vip3A toxins which contribute significantly to the overall insecticidal activity of the *B. thuringiensis* strains (11). This type of insecticidal toxin was first discovered in the culture supernatant of the *B. thuringiensis* strain AB88 and its production was detected from the vegetative growth phase to the end of the stationary phase and sporulation (12). Based on this atypical production profile, the authors proposed to name this new protein Vip, for Vegetative insecticidal protein. Since this discovery, it has been shown that the *vip3A* gene is present in about 50% of the *B. thuringiensis* strains tested and is located on large plasmids also harboring *cry1I* and *cry2A* genes (13, 14). Complete sequencing of the *B. thuringiensis* strain *kurstaki* HD1 indicates that the *vip3A* gene is located on the large plasmid pBMB299 which also carries the *cry1Aa, cry1Ia, cry2Aa* and *cry2Ab* genes (15).

The Vip3A toxins are specifically toxic against lepidopteran insects belonging to the Noctuidae family, including important agricultural pests like *Spodoptera exigua* and *Spodoptera frugiperda*, which are poorly susceptible to the Cry toxins (16, 17). Moreover, the specific receptors recognized by Vip3A toxins in the insect midgut are different from those recognized by the Cry toxins (18–20). These properties mean that *vip3A* genes are very often used in combination with *cry* genes in plant transgenesis pyramid strategies to increase plant resistance to insect pests but also to bypass the resistance to Cry toxins acquired by certain insects (21).

Despite the atypical expression pattern of *vip3A* genes and the importance they may play in the pathogenicity of *B. thuringiensis*, no comprehensive study of the regulation of their expression has ever been conducted. In this paper, we describe for the first time the expression kinetics of a *vip3A* gene, we identify its promoter and characterize a novel regulator that positively controls *vip3A* gene transcription during the stationary phase.

## Results

### Expression of the *vip3A* gene requires the presence of the plasmid pBMB299

To study the regulation of *vip3A* gene expression we used the *B. thuringiensis kurstaki* HD73 strain as a heterologous and naive host which does not carry naturally the *vip3A* gene (22). Specifically, we have used a *kurstaki* strain designated HD73^-^ which was cured of the plasmid pHT73 carrying the *cry1Ac* gene (23). The transcriptional activity of the DNA fragment located upstream from the *vip3A* gene present on the pBMB299 plasmid (NZ_CP004876.1) of the *B. thuringiensis kurstaki* HD1 strain was analyzed. This 709-bp DNA fragment was fused to the *lacZ* gene in the pHT304.18Z, a plasmid designed to study gene expression using promoter-*lacZ* transcriptional fusion in *B. thuringiensis* (Table 1) (24), and the HD73^-^ strain was transformed with the resulting plasmid pHT-P*_vip3_* (Fig. S1A). HD73^-^ (pHT-P*_vip3_*) bacteria were isolated on LB plates containing X-gal (50 μg/mL) and the plates were observed after 24 h of growth at 37°C. No blue colonies were observed, indicating that the bacterial cells did not produce β-galactosidase activity (Fig. S1B). This result suggested that a specific regulator required for *vip3A* expression was absent from the HD73^-^ strain while it was present in the parental HD1 strain. We hypothesized that this regulator was encoded by a gene of the pBMB299. To test this hypothesis we transferred the plasmid into the HD73^-^ Sm^R^ strain. *In silico* analysis of genes encoded on the pBMB299 revealed that a putative conjugal transfer protein (TraG) was present and suggested that this plasmid was conjugative. However, to compensate for the absence of a pBMB299-specific selection marker to select clones having received the plasmid, we developed a strategy to select a co-conjugation event between the pBMB299 and the mobilizable pHT1618K that confers a resistance to kanamycin. Indeed, it has been shown previously that the Gram-positive pBC16 replicon constituting pHT1618 is mobilizable by conjugation between *B. thuringiensis* strains (25). We introduced the pHT1618K (Table1) into the HD1 strain by electroporation and the HD1 (pHT1618K) Km^R^ was used as the donor to perform conjugation with the recipient strain HD73^-^Sm^R^. HD73^-^ Sm^R^ Km^R^ exconjugant clones having received the pHT1618K by a mobilization process were screened for the presence of the pBMB299 by PCR using the primer pair vip3-fw/vip3-rev targeting the *vip3A* gene. 14% of these exconjugants harbored the pBMB299 and one clone HD73^-^ Sm^R^ Km^R^ (pBMB299, pHT1618K) was selected. In agreement with the transfer of the pBMB299, the exconjugant clone produced bipyramidal crystals when grown on HCT plates for 4 days at 30°C (Fig. S2). The strain was then cured of the pHT1618K as previously described (26).

**Table 1:**
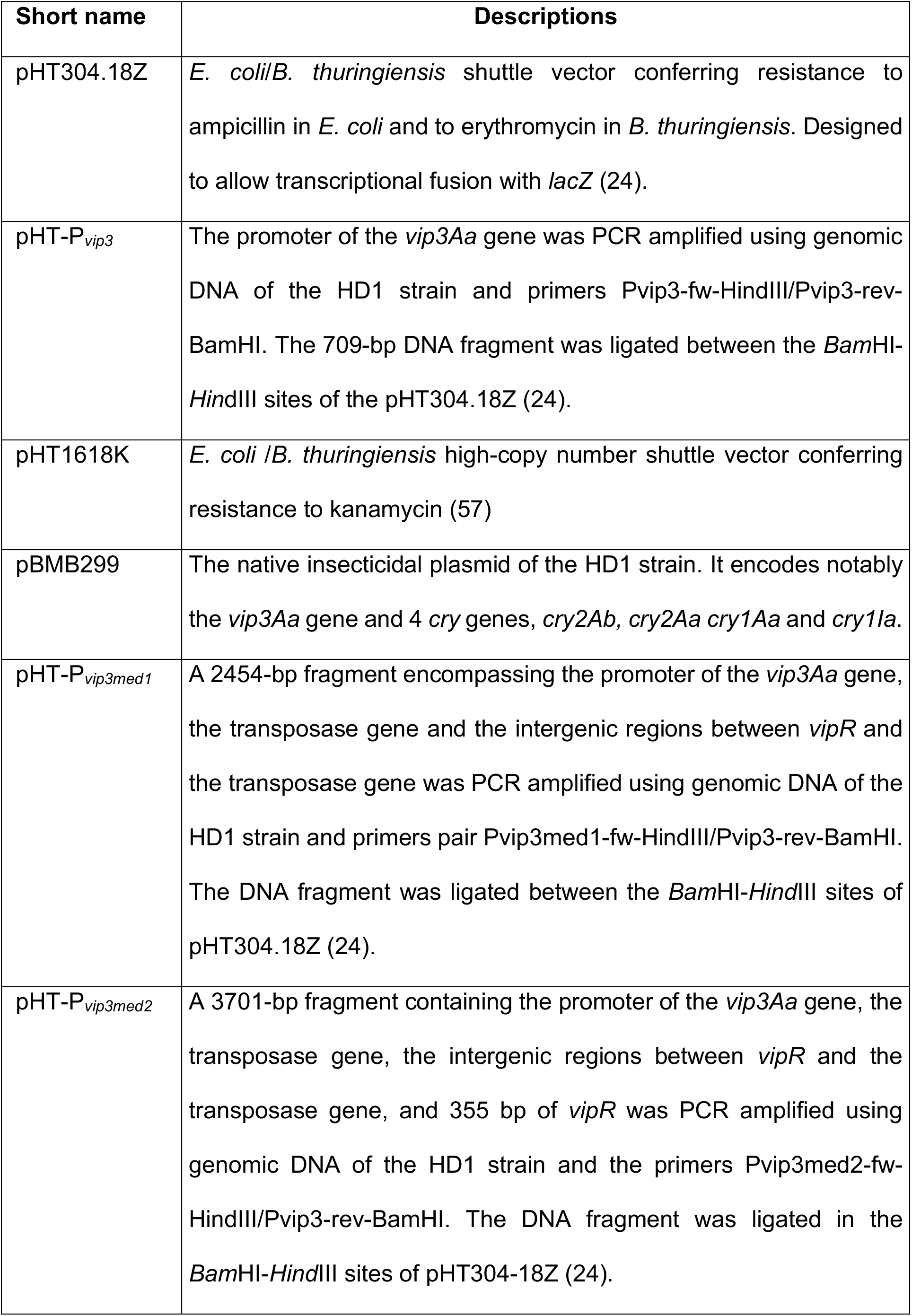
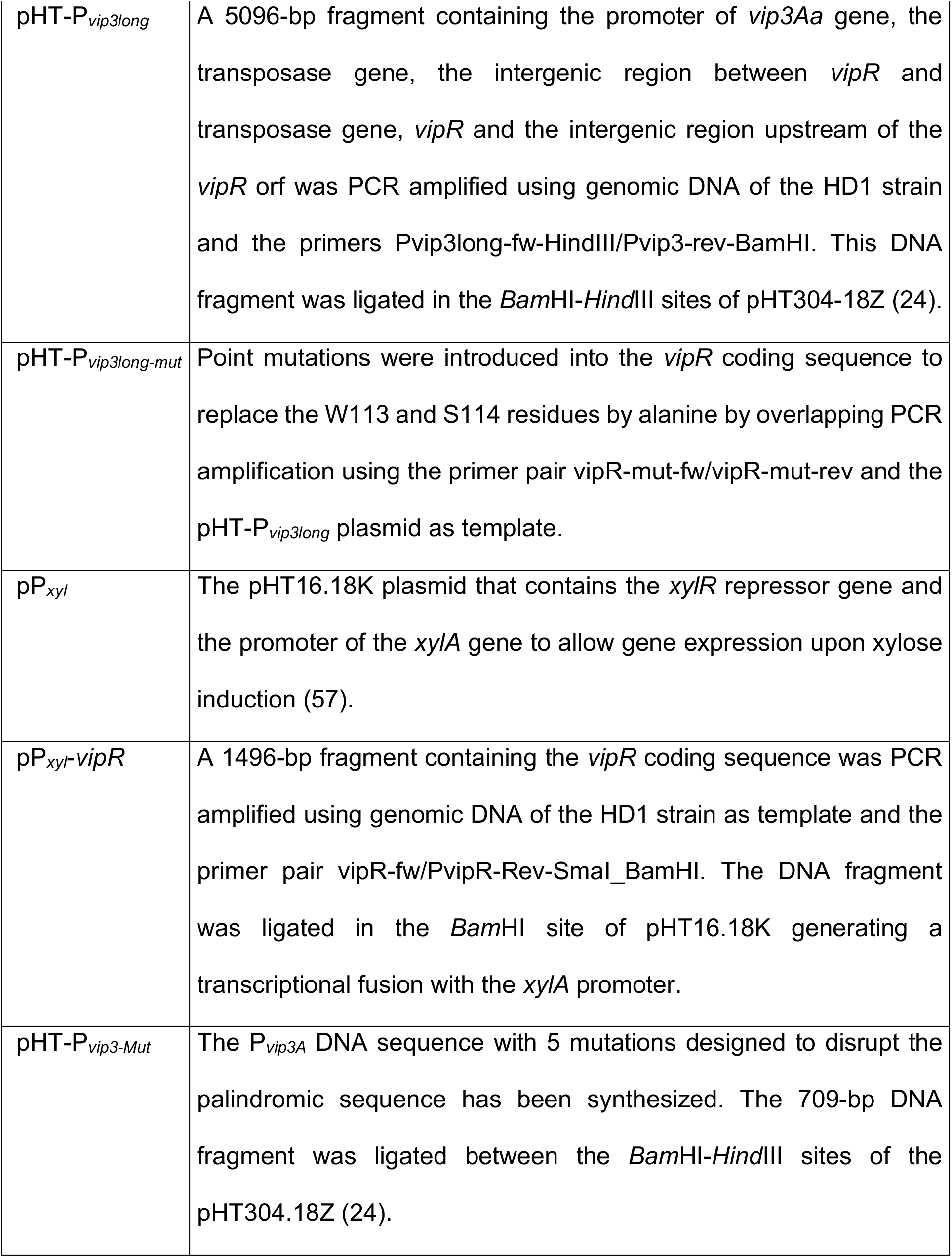
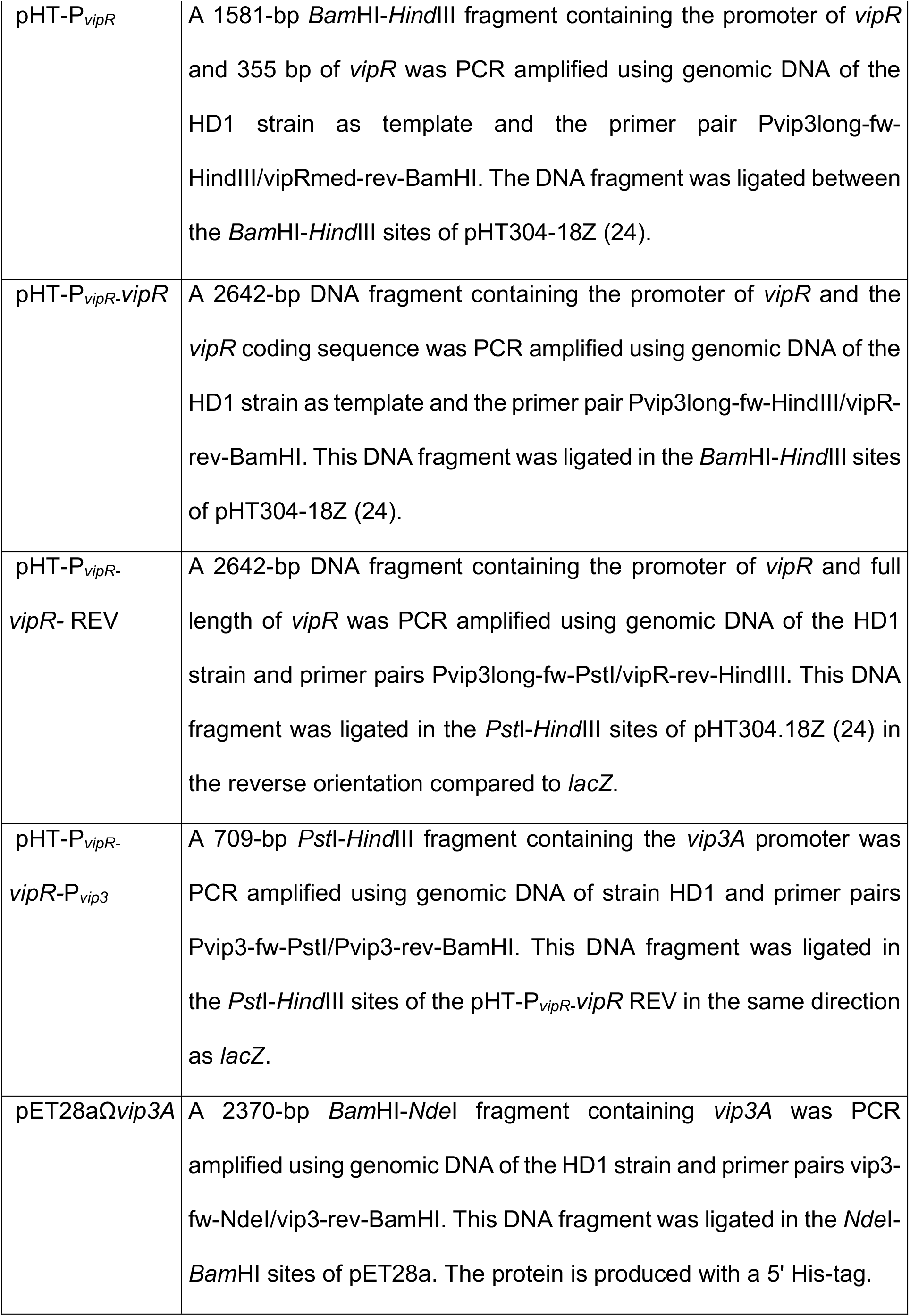
Plasmids used in the study.

We then analyzed the production of the Vip3A protein by Western blot using an anti-Vip3A11 antibody. The HD73^-^ Sm^R^ (pBMB299) and the HD73^-^ strains were cultivated in LB at 37°C, and bacterial cells and culture supernatants were collected at T1 and T4. The anti-Vip3A antibody did not reveal Vip3A in samples produced from the HD73^-^ Sm^R^ strain (Fig. 1). However, the Vip3A protein was detected in both the supernatant and the cell protein extract of the HD73^-^ Sm^R^ (pBMB299) strain. These results indicate that the HD73 strain is able to produce and secrete the toxin when it carries the pBMB299 and that the plasmid itself is able to specify the synthesis of Vip3A in the strain HD73. They therefore suggest that a regulator is encoded by a gene present on the pBMB299.

**Fig. 1.**
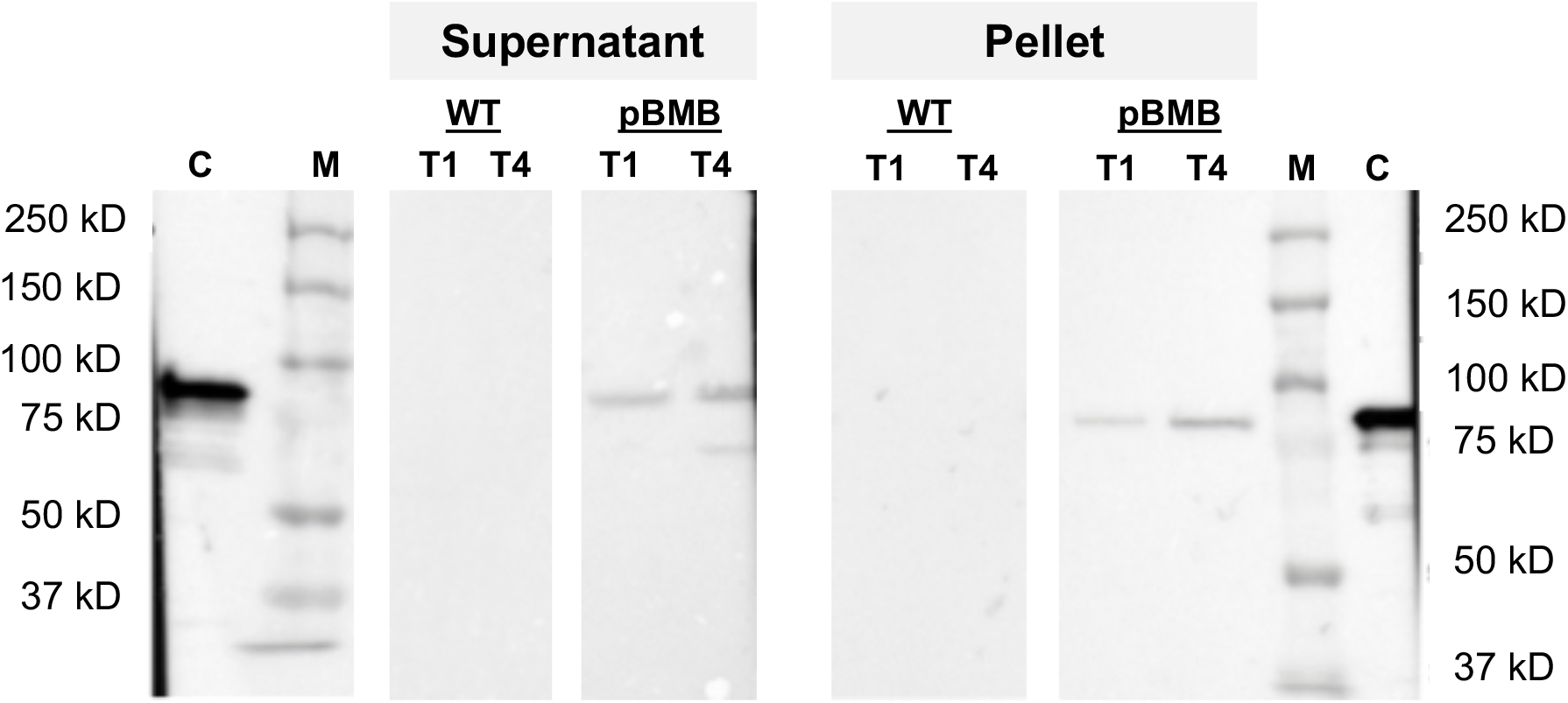
Expression of *vip3A* in the *B. thuringiensis kurstaki* HD73 Cry^-^ strain. Western blot analysis of Vip3Aa production in the *B. thuringiensis* HD73^-^ -WT- and HD73^-^ Sm^R^ (pBMB299) -pBMB-strains. Strains were cultured in LB medium at 37°C. Samples were collected 1h (T1) and 4h (T4) after the entry into stationary phase. The supernatant and cell pellet proteins were prepared as described in M&M. 20 μg of proteins were loaded in each well. 0,1 μg of purified Vip3Aa was used as a control. C – control, M – protein molecular weight marker.

### Identification of a gene involved in *vip3A* expression

To identify the pBMB299 gene(s) involved in *vip3A* expression, we performed a bioinformatic analysis on the *vip3A*-bearing plasmids from different *B. thuringiensis* strains (HD1, BGSC4C1, CT-43, HD12, HD29, IS5056, L7601, YBT-1520, YC-10). All of those are large plasmids with sizes ranging from 267 to 299 kb. In the HD1 strain, *vip3A* is located in a genetic environment containing multiple insertion sequences or transposase pseudogenes. Notably, *vip3A* is surrounded by two truncated DNA sequences showing similarities with the *tnpB* gene of the IS*1341* family. An IS*232* mobile element and a gene encoding a protein containing an N-terminal helix-turn-helix (HTH) domain putatively involved in DNA binding and annotated as a transcriptional antiterminator, are located upstream from *vip3A* (Fig. 2A). The *orf-HTH* gene is present at the same location in all the nine *B. thuringiensis* strains studied and the gene sequences displayed 90-100% of identity between strains (not shown). This gene was previously identified through a genomic analysis of the *B. thuringiensis* strain YBT1520 (15). The structural prediction of the protein encoded by the *orf-HTH* gene using the Phyre2 software (27) and an HHpred analysis revealed similarities to the *Bacillus anthracis* virulence regulator AtxA and the *Streptococcus pneumonia* virulence regulator Mga despite a low percentage of sequence identity (17 and 19 %, respectively). The presence of this HTH-containing protein gene in the vicinity of *vip3A* in all strains led us to investigate its role in the activation of the transcription of the *vip3A* promoter (P*_vip3_*). We constructed different transcriptional fusions with *lacZ* using DNA fragments of different lengths containing the P*_vip3_* and extending upstream to the end of the IS*232* ATPase coding sequence as schematized in Fig. 2A. The plasmids pHT-P*_vip3mid1_*, pHT-P*_vip3med2_* and pHT-P*_vip3long_* carrying these transcriptional fusions were transformed in strain HD73^-^. The resulting strains were plated on LB plates containing X-gal (Fig. S1C). The ß-galactosidase activity (blue color) was only detected in the colonies of the HD73^-^ (pHT-P*_vip3long_*) strain, suggesting that the DNA fragment containing the *orf*-HTH coding sequence was required for producing ß-galactosidase. These results were confirmed by determining the ß-galactosidase activity of the 4 transcriptional fusions P*_vip3_-lacZ*, P*_vip3mid1_-lacZ*, P*_vip3med2_-lacZ* and P*_vip3long_-lacZ* (Fig. 2B). The strains HD73^-^ (pHT-P*_vip3_*), HD73^-^ (pHT-P*_vip3mid1_*) and HD73^-^ (pHT-P*_vip3mid2_*) produced a very low amount of ß-galactosidase throughout the growth of the bacteria (Fig. 2B). On the contrary, the ß-galactosidase activity of the strain carrying the pHT-P*_vip3long_* was high during the vegetative growth and increased significantly 1 h after the onset of the stationary phase. These results suggest that the *orf-HTH* gene is involved in the expression of the *vip3A* gene.

**Fig. 2.**
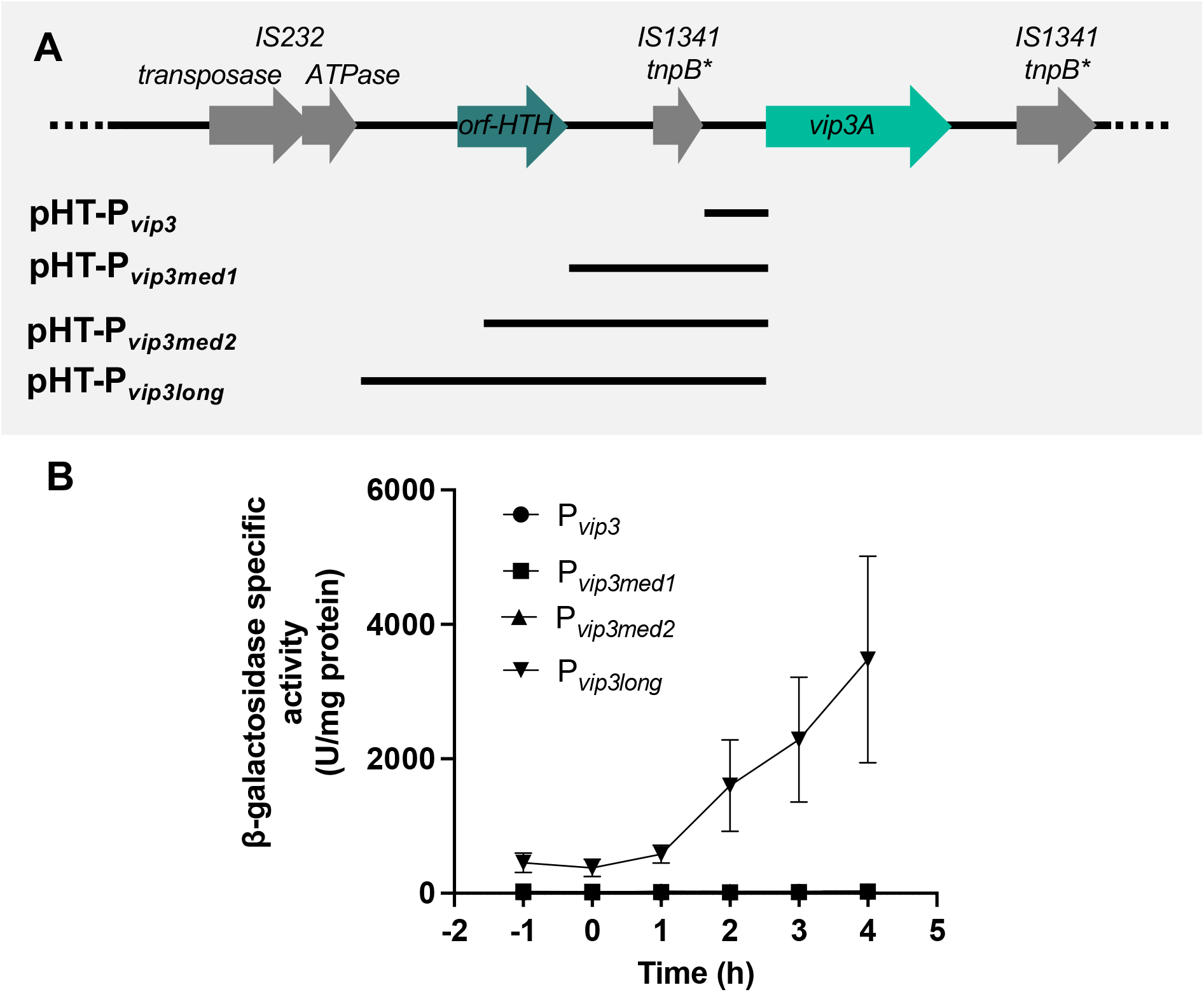
Characterization of the DNA region required for *vip3A* gene expression. A- Schematics representing the genes located upstream and downstream of the *vip3A* ORF and the DNA fragments screened for their ability to produce a transcriptional activity. The asterisk indicates a gene containing nonsense mutations. B- The ß-galactosidase activity of *B. thuringiensis* HD73^-^ strains harboring the pHT-P*_vip3_*, pHT-P*_vip3med1_*, pHT-P*_vip3med2_* or the pHT-P*_vip3long_* plasmid. The activity was assayed when *B. thuringiensis* HD73^-^ strains were grown in LB at 37°C. Time 0 corresponds to the entry of the bacteria into stationary phase. Data are the mean ± SEM, n=3.

### Analysis of the *vip3A* promoter region

We determined the 5’-end of the *vip3A* transcript using total RNA samples from HD1 cells harvested at T2. A putative *vip3A* transcriptional start site (TSS) was identified 403 bp upstream from the *vip3A* start codon (Fig. 3). This nucleotide is preceded by a potential −10 box (TATAAT) but no canonical SigA −35 box could be identified. Instead, analysis of the P*_vip3_* DNA sequence using the mFold software (28) found a palindromic DNA sequence having the potential to form a stem-loop structure ending 29 bp upstream of the putative TSS (Fig. 3). These data and the ß-galactosidase activity obtained with the P*_vip3long_-lacZ* transcriptional fusion (Fig. 2B) is consistent with the hypothesis that the protein encoded by the *orf-HTH* gene is the activator of the P*_vip3A_* promoter through its binding to the palindromic sequence.

**Fig. 3.**
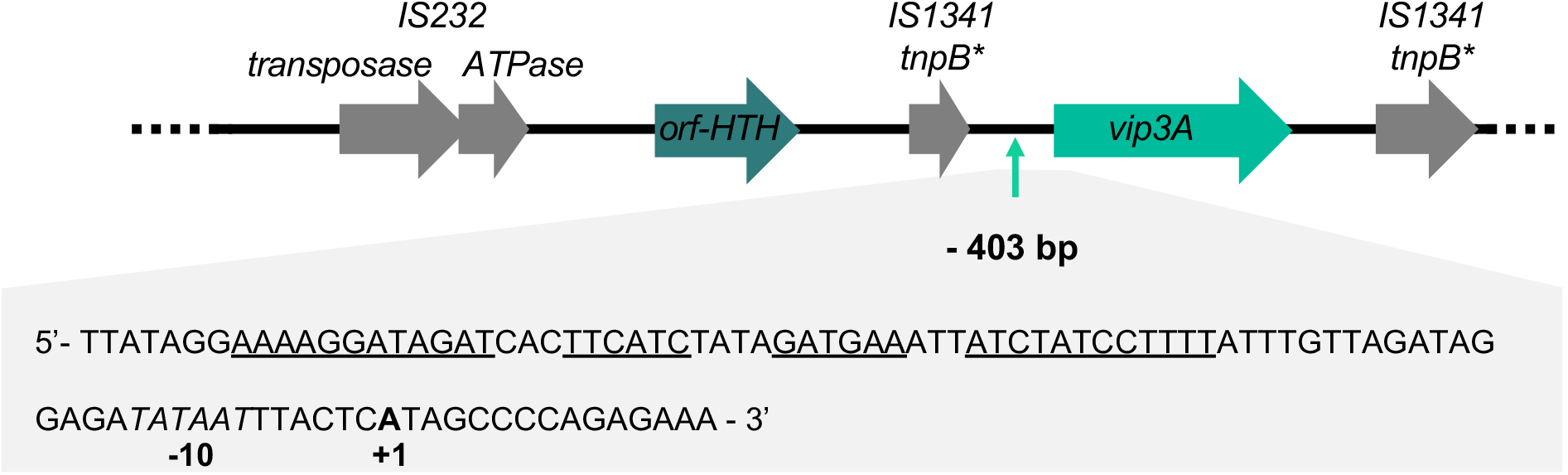
Characterization of the *vip3A* promoter elements. Schematic representation of the *vip3A* locus. The putative transcriptional start site is indicated with an arrow at position −403 relative to the *vip3A* start codon. The asterisk indicates a gene containing nonsense mutations. A focus on the DNA sequence that contains the *vip3A* promoter elements is given below. The mRNA 5’-end identified using RACE-PCR is indicated in bold. The DNA sequence corresponding to the putative −10-box is italicized. The palindromic sequences that are predicted to form a hairpin structure by the mFold software are underlined. The RNA used for the RACE-PCR was prepared from *B. thuringiensis* HD1 cells grown in LB medium and collected 2 hours after the entry into stationary phase.

### Characterization of the regulator of *vip3A* expression

The structure prediction of the protein encoded by the *orf-HTH* gene indicated a similarity with Mga, the virulence regulator of *S. pneumonia*. The alanine replacement of two amino acids in the HTH domain of Mga abolished its DNA-binding activity and regulation of its target genes (29). To determine whether the protein encoded by the *orf-HTH* gene is responsible for the activation of the *vip3A* promoter, its HTH domain was modified based on its alignment with the HTH domain of the protein Mga (Fig. S3). Two residues of the potential HTH domain, W113 and S114 were replaced by two alanines (Fig. 4A). Modeling of the mutated protein using the Phyre2 web server did not indicate a major structural change compared to the predicted 3D-structure of the orf-HTH protein. A transcriptional fusion between the P*_vip3long_* fragment containing the mutations and the *lacZ* gene was constructed in the pHT304.18Z plasmid and the HD73^-^ strain was transformed with the resulting pHT-P*_vip3long-mut_* plasmid to generate the HD73^-^ (pHT-P*_vip3long-mut_*) strain. Comparison of the β-galactosidase activity of the HD73^-^ (pHT-P*_vip3long-mut_*) and HD73^-^ (pHT-P*_vp3long_*) strains showed that the mutation of the two amino acids completely abolished the transcriptional activity of the P*_vip3long_* fragment (Fig. 4B). This result suggests that the protein encoded by the *orf-HTH* gene is involved in the activation of the P*_vip3A_* promoter during early stationary phase. This protein was named VipR for <u>Vip R</u>egulator.

**Fig. 4.**
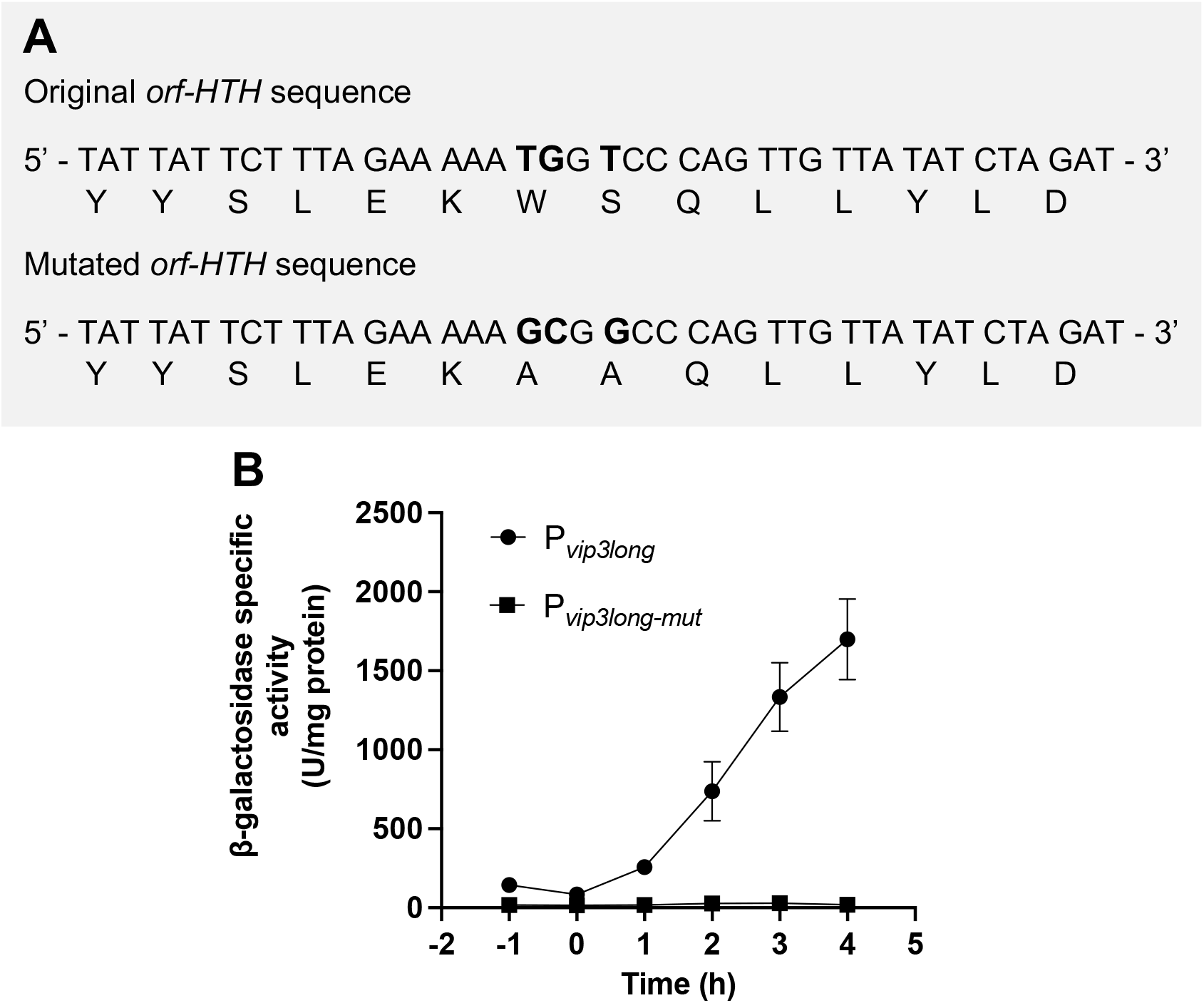
Mutations in the *orf-HTH* gene abolish the P*_vip3long_* transcriptional activity. A- Codons specifying the amino acids W113 and S114 were modified to each code for an alanine. Mutated bases are indicated in bold. B- ß-galactosidase activity of the HD73^-^ strains carrying the pHT-P*_vip3long_* or the pHT-P*_vip3long-mut_* plasmid. Bacteria were grown in LB at 37°C. Time 0 corresponds to the entry of the bacteria into stationary phase. Data are the mean ± SEM, n=4.

To confirm the activation of *vip3A* expression by VipR and to determine if the *lacZ* transcription produced with the P*_vip3long_-lacZ* transcriptional fusion originated from the promoter P*_vip3_*, we introduced the pHT-P*_vip3_* in the HD73^-^ that harbors the pP*_xyl_-vipR* plasmid. In this strain, the expression of *vipR* was directed from the xylose-inducible promoter P_xyl_ (Fig. 5A). The resulting HD73^-^ (pHT-P*_vip3_*, pP*_xyl_-vipR*) strain was cultivated in LB in the presence or absence of xylose in the culture and the ß-galactosidase activity of the cells was measured throughout the growth from T-1 to T7. We observed that the addition of xylose in the culture induced a strong increase in ß-galactosidase production in the stationary phase (Fig. 5B). This result demonstrated that the production of VipR activated *lacZ* transcription from the P*_vip3_*.

**Fig. 5.**
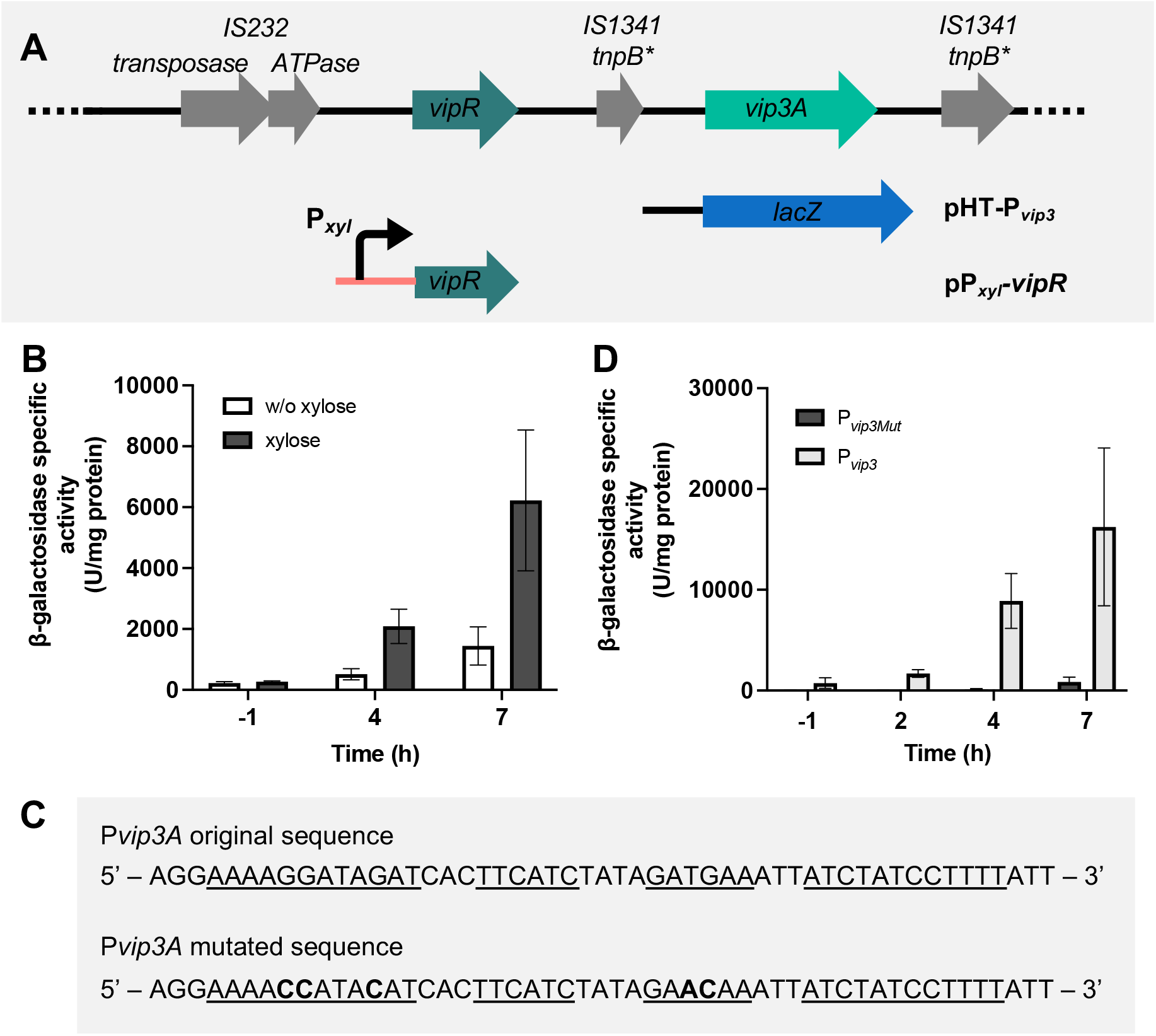
Activation of the promoter of *vip3* by VipR. **A-** Schematic representation of the constructs used to study the regulation of the *vip3* gene. The asterisk indicates a gene containing nonsense mutations. **B-** ß-galactosidase activity of the Bt HD73^-^ (pHT-P*_vip3_*, pP*_xyl_-vipR*) cells grown in the absence or in the presence of xylose (20 mM). Bacteria were cultured in LB at 37°C. **C-** DNA sequence of the *vip3A* promoter. Bases forming the palindrome are underlined. Mutated bases are indicated in bold. **D-** ß-galactosidase activity of the Bt HD73^-^ (pHT-P*_vip3_*, pP*_xyl_-vipR*) and HD73^-^ (pHT-P*_vip3-mut_*, pP*_xyl_-vipR*) cells grown in the presence of xylose (20 mM). Bacteria were cultured in LB at 30°C. Time 0 corresponds to the entry of the bacteria into the stationary phase. Xylose was added at T-1. Data are the mean ± SEM, n=3.

To determine whether the palindromic sequence located upstream of the putative *vip3A* TSS (Fig. 3) was involved in the control of *vip3A* expression, mutations were introduced to disrupt the palindrome and eventually to prevent this DNA sequence to form a stem-loop structure (Fig. 5C). A transcriptional fusion between the mutated P*_vip3_* promoter and *lacZ* was created in the pHT304.18Z and the resulting plasmid pHT-P*_vip3-mut_* was introduced in the strain HD73^-^ (pP*_xyl_-vipR*). Measurement of the ß-galactosidase activity of the strains HD73^-^ (pP*_xyl_-vipR*, pHT-P*_vip3_*) and HD73^-^ (pP*_xyl_-vipR*, pHT-P*_vip3-mut_*) showed that the transcriptional activity of the mutated promoter was drastically lower than that of the wild-type promoter (Fig. 5D). This result indicates that the integrity of the palindromic sequence is required for full activation of *vip3A* transcription by VipR, possibly by forming a stem-loop structure.

### The *vipR* gene is autoregulated

The expression kinetics of *vipR* was studied using a 1581-bp DNA fragment (P*_vipR_*) corresponding to the intergenic region between *IS232 ATPase* and the first 355 bp of the *vipR* coding sequences (Fig. 6A). This DNA fragment was transcriptionally fused with *lacZ* in the pHT304.18Z plasmid and the HD73^-^ strain was transformed with the resulting pHT-P*_vipR_* plasmid. ß-galactosidase activity of strain HD73^-^ (pHT-P*_vipR_*) was measured throughout the growth of bacteria cultivated in LB medium (Fig. 6B). Results showed that P*_vipR_* expression was low during the exponential phase of growth and increased 4-fold from T0 to T3. To determine if *vipR* expression is autoregulated, we introduced the plasmid pHT-P*_vipR_* in the strains HD73^-^ (pP*_xyl_*) and HD73^-^ (pP*_xyl_-vipR*) and compared the ß-galactosidase activity of the two strains (Fig. 6C). The results showed that *vipR* transcription is significantly increased when VipR is produced in the bacterial cells under the control of P*_xyl_*. Therefore, VipR presents the characteristics of an autoregulated transcriptional activator.

**Fig. 6.**
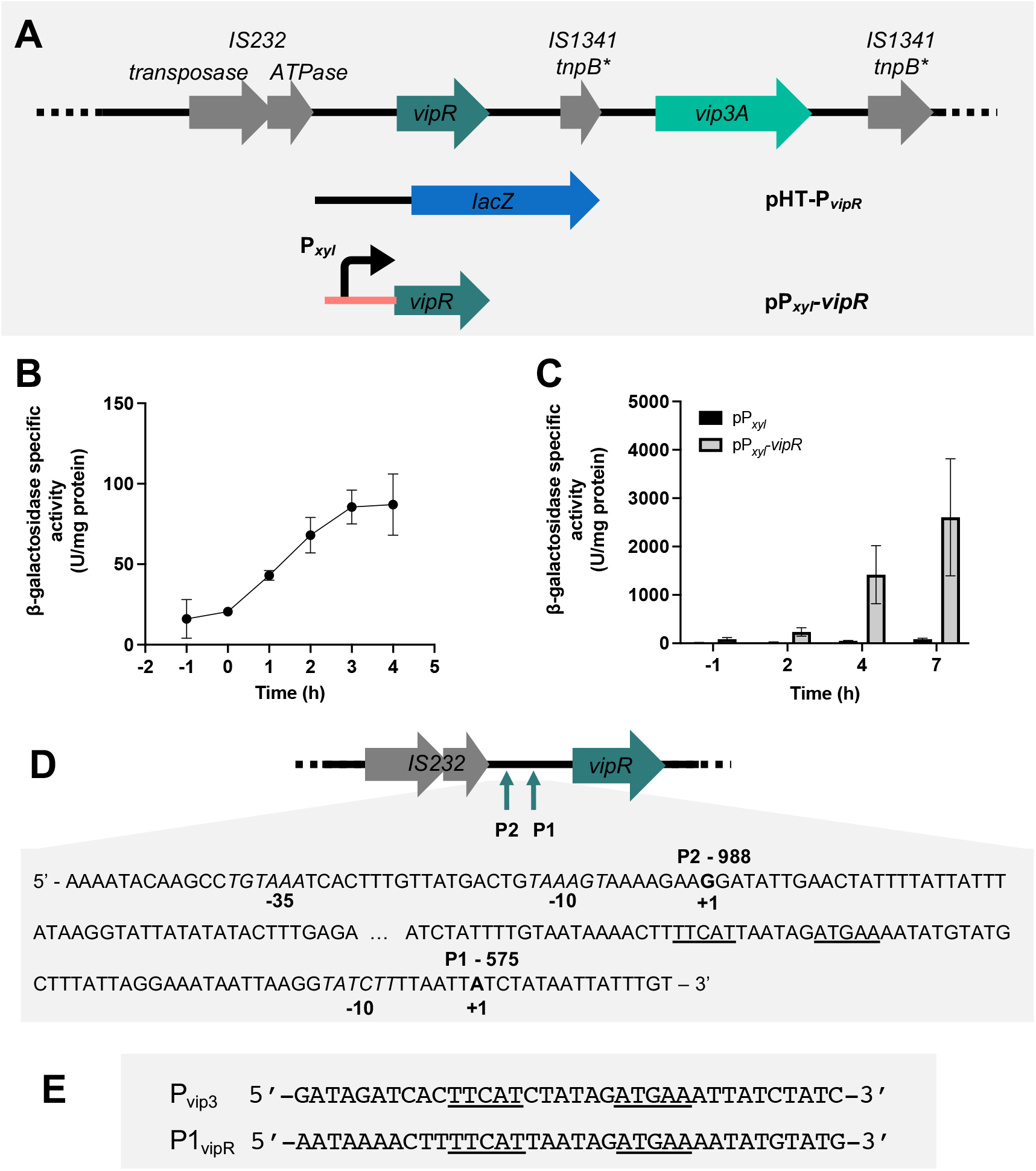
VipR is an autoregulated transcriptional activator. **A-** Schematic representation of the constructs used to study the regulation of the *vipR* gene. **B-** ß-galactosidase activity of the Bt HD73^-^ (pHT-P*_vipR_*) cells grown in LB at 37°C. **C-** ß-galactosidase activity of the Bt HD73^-^ (pHT-P*_vipR_*, pP*_xyl_-vipR*) and Bt HD73^-^ (pHT-P*_vipR_*, pP*_xyl_*) cells grown in the presence of xylose (20 mM) at 30°C. Time 0 corresponds to the entry of the bacteria into the stationary phase. Xylose was added at T-1. Data are the mean ± SEM, n=3 **D-** Schematic representation of the *vipR* genetic organization. The putative transcriptional start sites are indicated with an arrow. P1 and P2 are located at position - 575 and −988 to the *vip3A* start codon, respectively. A focus on the DNA sequence that contains the *vipR* promoter elements is given. The 5’ ends of the two *vipR* mRNA identified using RACE-PCR are indicated in bold. The DNA sequence corresponding to the putative – 10-boxes are italicised. The palindromic sequence in P1 is underlined. The RNA used for the RACE-PCR was prepared from Bt HD73^-^ pHT-P*_vip3long_* cells grown in LB medium and collected at T2. **E-** Alignment of the DNA sequences of the *vipR* and *vip3A* promoters highlighting the conserved palindromic sequences.

We then determined the 5’-end of the *vipR* transcript using total RNA samples from HD73^-^ (pHT-P*_vip3long_*) bacteria harvested at T2. A proximal *vipR* putative TSS named P1 was identified 575 bp upstream from the *vipR* start codon (Fig. 6D). This putative start is preceded by a potential −10 box (TATCTT). As for *vip3A*, no canonical SigA - 35 box could be identified but a 16-bp palindromic sequence is present 32 bp upstream of this putative TSS. Alignment of this sequence with the palindrome sequence of the *vip3A* promoter allowed us to identify a conserved DNA sequence that might be the target of the regulation by VipR (Fig. 6E) and suggests that P1 is the autoregulated *vipR* promoter. The 5’-RACE analysis of the *vipR* promoter indicated another 5’-end 998 bp upstream of the *vipR* start codon. This distal putative TSS, named P2, is preceded by a canonical −10 box (TAAAGT). A putative SigA −35 box (TGTAAAA) is correctly positioned 17 bp upstream of the −10-box suggesting that *vipR* transcription from the P2 promoter would be SigA-dependent. Identification of two possible *vipR* promoters is consistent with the results obtained with a Northern blot analysis of *vipR* transcription showing the detection of two transcripts in RNA samples of the HD73^-^ (pHT-P*_vip3long_*) strain (Fig. S4). Discrete bands corresponding to the two transcripts were not detected in the HD1 and HD73^-^ (pBMB299) RNA samples. However, a light smear at the same place was present. The higher abundancy of the *vipR* transcripts in the HD73^-^ (pHT-P*_vip3long_*) strain may be due to a higher copy number of the pHT-P*_vip3long_* compared to the pBMB299.

### Identification of putative VipR-regulated genes on the plasmid pBMB299

We searched for DNA sequences showing similarities with the putative VipR binding site in the plasmid pBMB299, and identified a total of 7 conserved sequences located in the 5’-untranslated region of coding sequences (Fig. 7). The relevance of these sequences as potential VipR binding sites is greatly reinforced by the presence of a putative SigA −10 box 17 bp downstream from the last nucleotide of the consensus sequence. This result strongly suggests that the transcription of these 7 genes is at least partly controlled by VipR. In addition to *vip3A* and *vipR*, it appears that the expression of the *cry1I* gene expression might also be controlled by VipR. This result would be consistent with the observation that Cry1I toxin (formerly CryV) was produced in early stationary phase and exported, like Vip3A (30). More surprisingly, the two *cry2A* genes (formerly *cryB* or *cryII*) are also located downstream from a putative VipR box. These genes are known to be transcribed by sporulation-specific sigma factors (31, 32). A VipR-dependent expression would mean that the Cry2A toxins are also produced prior to the sporulation process, and thus prior to the formation of the parasporal crystal inclusion. Finally, two genes encoding N-acetylmuramoyl-L-alanine amidases (designated Amidase 1 and 2 on figure 7) are also located downstream of a putative VipR box.

**Fig. 7.**
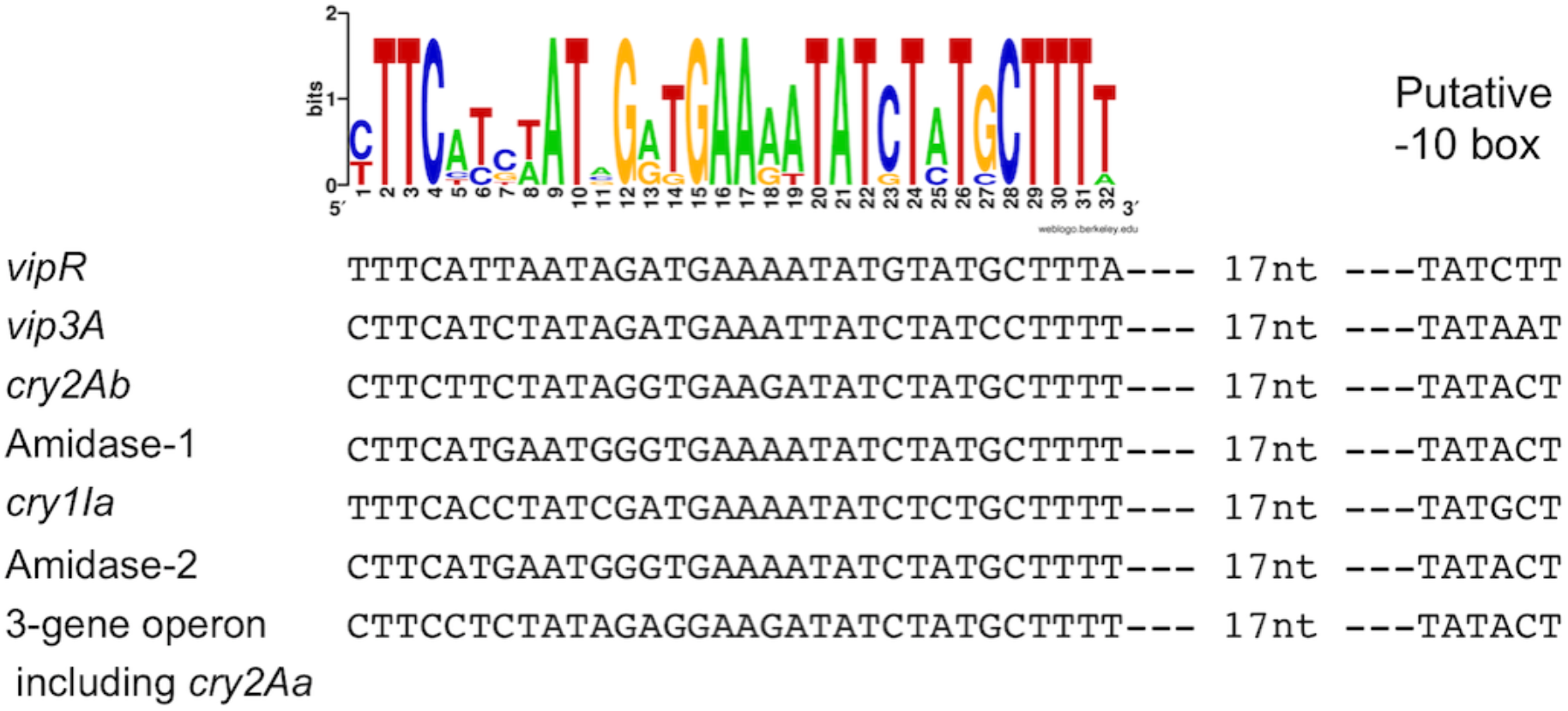
Alignment of the conserved sequences found in the pBM299 plasmid. The name of the gene putatively controlled by VipR is indicated on the left. Distance between the putative −10-box and the last nucleotide of the conserved motif is indicated. A consensus is shown on top as a sequence logo in which the height of the letters in bits is proportional to their frequency.

### *vip3A* and *vipR* expression is strongly increased in a Δ*spo0A* genetic background

The activation of *vipR* and *vip3A* expression at the onset of stationary phase led us to address the role of a key regulator of the transition phase in Bacilli, Spo0A (33). *vipR* expression in the *spo0A* mutant was studied using a 2644-bp DNA fragment that includes the 5’-untranslated region upstream from *vipR* and the *vipR* coding sequence. This DNA fragment was cloned upstream of the *lacZ* gene in pHT304.18Z and the resulting plasmid, pHT-P*_vipR_-vipR* (Fig. 8A), was introduced in the HD73^-^ and HD73^-^ Δ*spo0A* strains. *vipR* transcription was compared in these two genetic backgrounds. Levels of ß-galactosidase produced by the bacteria indicated that, in the presence of *vipR*, a 40-fold increase in *vipR* transcription was observed in the Δ*spo0A* mutant (Fig. 8B).

**Fig. 8.**
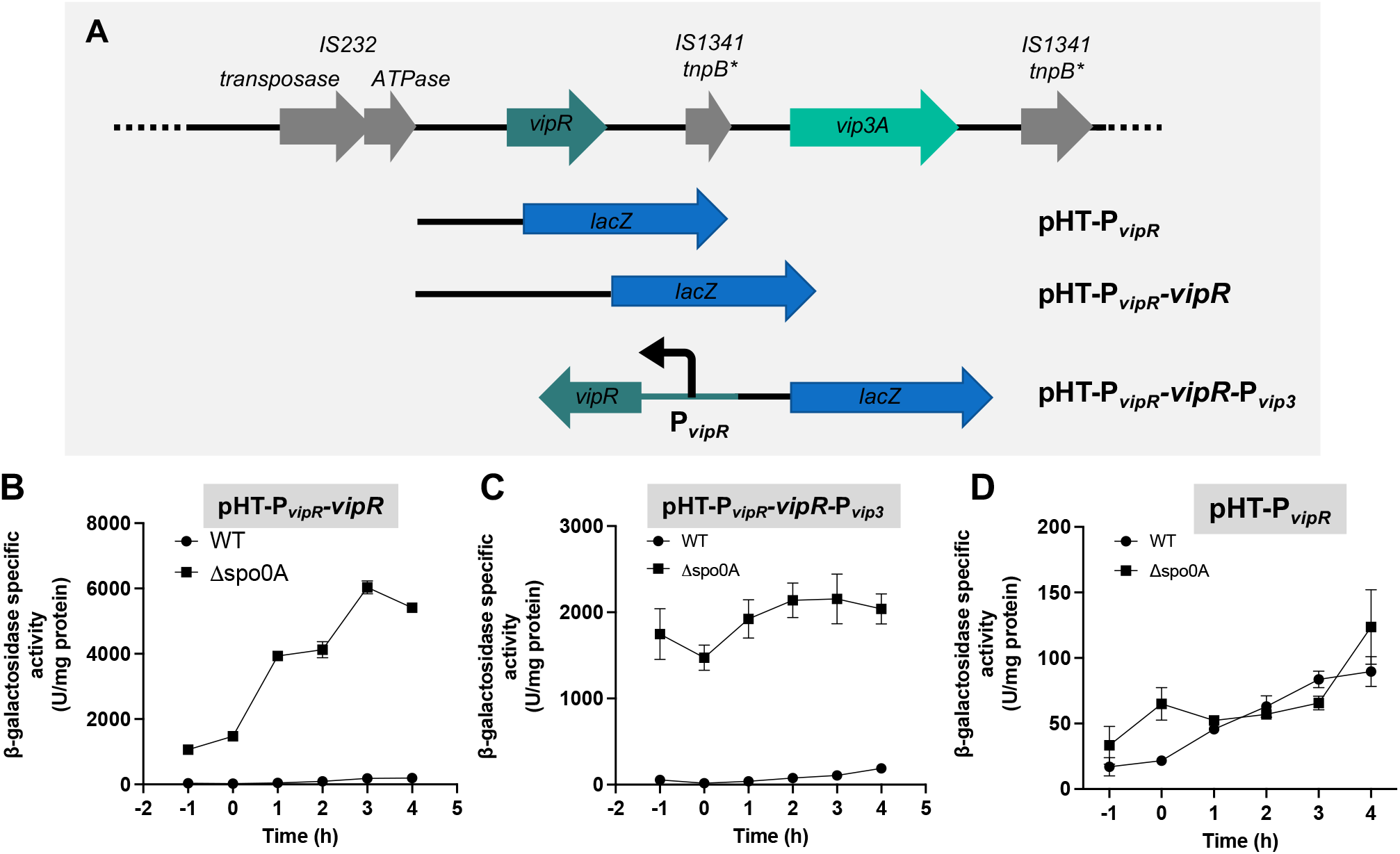
Expression of *vip3A* is increased in the Bt HD73 Cry^-^ Spo0A^-^. **A-** Schematic representation of the constructs used to study the regulation of the *vip3* and *vipR* genes in the sporulation mutant strain. **B-** ß-galactosidase activity of the Bt HD73^-^ (pHT-P*_vipR_-vipR*) and Bt HD73^-^ Spo0A^-^ (pHT-P*_vipR_-vipR*) cells. Data are the mean ± SEM, n= at least 3. **C-** ß-galactosidase activity of the Bt HD73^-^ (pHT-P*_vipR_-vipR*-P*_vip3_*) cells and Bt HD73^-^ Spo0A^-^(pHT-P*_vipR_-vipR*-P*_vip3_*). Data are the mean ± SEM, n= at least 4. **D-** ß-galactosidase activity of the Bt HD73^-^ (pHT-P*_vipR_*) and Bt HD73^-^ Spo0A^-^(pHT-P*_vipR_*) cells. Strains were grown in LB at 37°C. Time 0 corresponds to the entry of the bacteria into the stationary phase. Data are the mean ± SEM, n= 3.

To determine whether the *vipR* transcriptional increase resulted in an increase in *vip3A* expression, we measured the activation of P*_vip3A_* by VipR in the HD73^-^ Δ*spo0A* mutant. In the P*_vip3long_* DNA sequence, the *vipR* and *vip3A* orfs are in the same direction and no terminator sequence has been identified downstream *vipR*. We therefore cannot rule out the possibility that *vipR* transcription generates a polycistronic mRNA that includes *vip3A*. Thus, in order to disconnect *vipR* transcription from P*_vip3A_* transcriptional activity, we constructed a plasmid carrying a *vipR* expression cassette oriented in the opposite direction compared to the P*_vip3A_-lacZ* transcriptional fusion. A 2642-bp DNA fragment that includes the 5’-untranslated region upstream from *vipR* and the *vipR* coding sequence was cloned in the reverse orientation upstream of the P_vip3_ DNA fragment in the pHT304.18Z plasmid (Fig. 8A). The HD73^-^ and HD73^-^ Δ*spo0A* strains were transformed with the resulting plasmid pHT-P*_vipR_-vipR*-P*_vip3_* and the ß-galactosidase activity of the two strains was compared (Fig. 8C). We observed that *vip3A* expression was strongly increased in the HD73^-^ Δ*spo0A* strain compared to the HD73^-^ strain.

Bioinformatic analysis of the DNA sequence upstream of the *vipR* coding-sequence did not allow us to identify a DNA sequence corresponding to the *B. subtilis* Spo0A box (34). However, to determine if Spo0A controls *vipR* expression at a transcriptional level, the pHT-P*_vipR_* plasmid was introduced in the HD73^-^ Δ*spo0A* strain and the ß-galactosidase activity of the strain was compared to the activity of the HD73^-^ (pHT-P*_vipR_*) cells. The results showed that, in the absence of VipR, *vipR* transcription is similar in the two strains (Fig. 8D), indicating that Spo0A does not affect the transcriptional activity from the distal promoter P2.

## Discussion

Vip3A was previously identified as a major insecticidal toxin of *B. thuringiensis* produced and secreted during vegetative growth until late stationary phase (12). Using transcriptional fusions between the *vip3A* promoter and the *lacZ* reporter gene, we show that *vip3A* expression is significantly increased at the onset of stationary phase and continues for several hours at a high level when bacteria are grown in a rich medium such as LB. We characterized the transcriptional regulator VipR responsible for the activation of *vip3A* expression at the onset of stationary phase. The *vipR* gene is located about 2.5 kb upstream from the *vip3A* gene on the pBMB299 plasmid and the *vipR-vip3A* locus is nearly identical in all *B. thuringiensis* strains harboring these genes (Table S1). Therefore, we assume that the results we describe in this study should be valid for all these *B. thuringiensis* strains. This claim agrees with a result showing that in *B. thuringiensis* strain KN11, Vip3A is specifically produced during the stationary phase and accumulates in the cytoplasm of bacteria grown in twofold diluted LB medium (35). The production of Vip3A during vegetative growth (12) and the weak *vip3A* expression we observed during the exponential growth phase (Fig. 2B and 4B) might be due to the weak VipR-independent transcription from the *vipR* promoter region (Fig. 6B). This hypothesis is supported by the fact that mutations in the HTH domain of VipR (Fig. 4B) or in the putative VipR box (Fig. 5D) abolished transcription from the P*_vip3A_* promoter throughout the bacterial culture. Moreover, a transcriptional fusion between the DNA fragment P*_vipR_* and the *lacZ* gene yielded a low but significant production of ß-galactosidase in the absence of VipR, thus reflecting a VipR-independent transcription from this promoter region (Fig 6B). The determination of the two putative transcription start sites upstream from *vipR* indeed reveals a putative SigA-dependent promoter (P2) which might be responsible for this low transcription.

The characterization of the main *vip3A* and *vipR* promoters indicates the presence of a putative −10 box resembling that recognized by the vegetative Sigma A factor of *Bacillus* (TATAAT) (36). However, the −35 region of both promoters does not match the consensus sequence of Sigma A promoters (TTGACA). The putative binding of VipR a few nucleotides upstream of the *vip3A* and *vipR* promoters may compensate for the lack of the −35 box by recruiting the RNA polymerase to the promoter, thereby allowing the initiation of transcription (37). Taken together these results suggest that VipR interacts with SigA-associated RNA polymerase to activate its own transcription and that of *vip3A* at the onset of stationary phase.

The positive autoregulation of VipR and the presence of a conserved DNA sequence in the promoters of *vip3A* and *vipR* suggest a positive regulatory loop specifically active during the stationary phase. Such a regulatory system requires two complementary conditions: i) a constitutive expression of the regulatory gene to trigger the activation of the regulatory loop, and ii) a transcriptional or post-transcriptional activation of the regulator at the beginning of the stationary phase. Constitutive expression of *vipR* may be due to an upstream promoter, and, as described above, such a transcriptional activity can indeed be detected from the distal promoter P2. This weak, constitutive VipR-independent transcription both explains the low production of Vip3A during bacterial exponential growth and the initiation of the regulatory loop activating *vipR* transcription. As for the specific activation of *vipR* expression at the onset of stationary phase, two main mechanisms may be involved. First, a mechanism involved at the transcriptional level and resulting in increased transcription of *vipR;* second, a post-transcriptional mechanism resulting in functional activation of the VipR regulator. Regulation at the transcriptional level may be due to global transition state transcriptional regulators like CodY, Spo0A, SinR, or AbrB (1, 38–41). We showed that the weak VipR-independent expression, presumably directed by the distal promoter P2, was constitutive and unaffected by a *spo0A* deletion (Fig. 8D), suggesting that activation of the *vipR-vip3A* locus during the stationary phase did not depend on the promoter P2. In contrast, the VipR-dependent expression of *vipR* was drastically increased in a Δ*spo0A* mutant indicating that Spo0A represses *vipR* transcription. Spo0A and the Spo0A box are well conserved among *Bacillus* (6, 34, 42, 43) and we did not find any putative Spo0A box in the *vipR* promoter region, suggesting that the effect of Spo0A on *vipR* expression was indirect. Since *vipR* expression depends on the activity of the VipR protein, this effect of the Δ*spo0A* mutation may be indirect through a derepression of *vipR* transcription by releasing the action of a repressor at the onset of stationary phase. Alternatively, the activity of VipR may depend on a functional activation.

VipR presents structural homologies with the *B. anthracis* AtxA and the *Streptococcus* Mga transcriptional regulators belonging to the PRD-Containing Virulence Regulators (PCVR) family (44). These regulators are characterized by the presence of two N-terminal HTH domains, two centralized phosphoenolpyruvates: carbohydrate phosphotransferase system-regulated domains (PRD) and a C-terminal EIIB-like domain. The PRD domains contain conserved phosphorylated histidine residues and are found in sugar operon regulators. The phosphorylation state of the AtxA PRD1 and PRD2 histidine residues regulates its activity and dimerization. Mutations affecting these residues result in altered level of toxin and capsule gene expression (45, 46). In *S. pyogenes*, Mga harbors two phosphohistidines in PRD1 and one phosphohistidine in PRD2 but only the PRD1 histidine residues have been implicated in protein function. Phosphoablative and phosphomimetic Mga PRD histidines mutant are attenuated in a group A *streptococcus* virulence model (47). Sequence analysis of the predicted PRDs of VipR indicates that a histidine residue is present at the location of the phosphohistidines H379 of AtxA and H324 of Mga. Two putative phosphohistidines (H191 and H215) are present in the PRD1 but they do not align with H199 of AtxA nor H204 or H270 of Mga. These elements suggest that VipR activity might be regulated by PTS-mediated phosphorylation and that Vip3A toxin expression may be linked to the bacterial metabolism.

While Cry proteins alone are highly toxic to susceptible insects, the presence of spores has a synergistic effect and significantly increases the insecticidal activity of Cry toxins (10, 48). This effect is due to a wide variety of virulence factors that make *B. thuringiensis* a true and highly effective entomopathogen (9). In strains, such as *kurstaki* HD1, Vip3A might have an important role in the pathogenicity. Indeed, Donovan and colleagues (11) have shown that deletion of the *vip3A* gene in the HD1 strain significantly decreased toxicity against lepidopteran insects specifically susceptible to Vip3A (eg. *S. exigua*), and their results suggested that this effect was due to the production of Vip3A during development of the bacteria in the insect gut. The determination of the expression profile of *vip3A* and the characterization of the VipR regulator provide new insights into the pathogenicity of *B. thuringiensis* in insects. An important issue to be verified is that VipR controls the expression of the other insecticidal toxin genes (*cry1I, cry2Aa* and *cry2Ab*) during the stationary phase, and subsequently it will be interesting to determine the role of all these factors during infection. In addition, it will be also interesting to determine the role of the two N-acetylmuramoyl-L-alanine amidases. In *B. thuringiensis*, it was shown that such an enzyme was involved in mother cell lysis at the end of the sporulation process (49), and in *Clostridium difficile* a peptidoglycan-degrading amidase allowed the release of toxins during stationary phase (50). Based on these functions and their suspected relation with VipR, it is tempting to hypothesize that the two amidases encoded by the plasmid pBMB299 are involved in the export of Vip3A and possibly Cry1I.

Overall, results regarding the pathogenicity of *B. thuringiensis* indicate that ingestion of Cry toxins causes toxemia followed by germination of bacterial spores and development of bacteria in the insect gut (1, 9). In contrast to a massive spraying of biopesticides, it is predictable that in the wild environment the amount of Cry toxins ingested by an insect is often not sufficient to kill it. Under these conditions, the production, in the insect gut, of additional toxins such as Vip3A could have an essential function in strengthening the infection and ensuring the multiplication of the bacteria. All these aspects deserve to be studied for a better understanding of the ecology of a bacterium used worldwide as a biopesticide and disseminated in large quantities in the environment.

## Materials and methods

### Strains and plasmids construction

The acrystalliferous strain *B. thuringiensis* HD73 Cry^-^ belonging to serotype 3 (22) was used as a heterologous host throughout this study and was designated as HD73^-^. *Escherichia coli* strain DH5α was used as the host strain for plasmid construction. *E. coli* strain BL21 λDE3 (Invitrogen) was used to produce the Vip3Aa protein. *E. coli* strain ET12567 (51) was used to prepare demethylated DNA prior to be used for transformation of *B. thuringiensis* by electroporation (52, 53). The plasmids and bacterial strains used in the study are listed in Table 1 and 2, respectively. Bacteria were routinely grown in LB medium at 37°C, and *B. thuringiensis* cells were cultured at 30°C when indicated. Time 0 was defined as the beginning of the transition phase between the exponential and stationary phases of *B. thuringiensis* growth. The following concentrations of antibiotics were used for *B. thuringiensis* selection: erythromycin (10 μg/mL), streptomycin (200 μg/mL) and kanamycin (200 μg/mL). For *E. coli* selection, kanamycin (20 μg/mL) and ampicillin (100 μg/mL) were used. When needed, xylose (20 mM) was added to the culture.

**Table 2:** Strains used in the study.

| Strain | Description | Antibiotic resistance |
| --- | --- | --- |
| HD73<sup>-</sup> | *B. thuringiensis kurstaki* HD73 cured of the plasmid pHT73 carrying the *cry1Ac* gene. This Cry<sup>-</sup> strain does not produce any crystal protein. | - |
| HD1 | *B. thuringiensis kurstaki* HD1 strain. The strain harbors the insecticidal plasmid pBMB299. | - |
| HD73<sup>-</sup> Sm<sup>R</sup> | Strain HD73<sup>-</sup> which is resistant to the streptomycin. This strain was used as the recipient strain for the pBMB299 and pHT1618K plasmids in the conjugation experiment. Lab stock. | Sm<sup>R</sup> |
| HD1<br>(pBMB299,<br>pHT1618K) | Strain HD1 transformed with the pHT1618K plasmid. This strain was used as the donor strain in the conjugation experiment. | Km<sup>R</sup> |
| HD73<sup>-</sup> Sm<sup>R</sup><br>(pBMB299,<br>pHT1618K) | Strain HD73<sup>-</sup> Sm<sup>R</sup> ex-conjugating. This strain received the pBMB299 and pHT1618K plasmids by conjugation from the HD1 strain. | Sm<sup>R</sup>, Km<sup>R</sup> |
| HD73<sup>-</sup> Sm<sup>R</sup><br>(pBMB299) | Strain HD73<sup>-</sup> Sm<sup>R</sup> harboring the pBMB299. The strain was cured from the pHT1618K plasmid. | Sm<sup>R</sup> |
| HD73<sup>-</sup><br>(pHT-P<sub>*vip3*</sub>) | Strain HD73<sup>-</sup> harboring the pHT-P<sub>*vip3*</sub> plasmid to measure the activity of the *vip3A* promoter. | Em<sup>R</sup> |
| HD73<sup>-</sup><br>(pHT-P<sub>*vip3med1*</sub>) | Strain HD73<sup>-</sup> harboring the pHT-P<sub>*vip3med1*</sub> plasmid to measure the transcriptional activity of the P<sub>*vip3med1*</sub> DNA fragment. | Em<sup>R</sup> |
| HD73<sup>-</sup><br>(pHT-P<sub>*vip3med2*</sub>) | Strain HD73<sup>-</sup> harboring the pHT-P<sub>*vip3med2*</sub> plasmid to measure the transcriptional activity of the P<sub>*vip3med2*</sub> DNA fragment. | Em<sup>R</sup> |
| HD73<sup>-</sup><br>(pHT-P<sub>*vip3long*</sub>) | Strain HD73<sup>-</sup> harboring the pHT-P<sub>*vip3long*</sub> plasmid to measure the transcriptional activity of the P<sub>*vip3long*</sub> DNA fragment. | Em<sup>R</sup> |
| HD73<sup>-</sup><br>(pHT-P<sub>*vip3long-mut*</sub>) | Strain HD73<sup>-</sup> harboring the pHT-P<sub>*vip3long-mut*</sub> plasmid to measure the transcriptional activity of the mutated P<sub>*vip3long-mut*</sub> DNA fragment. | Em<sup>R</sup> |
| HD73<sup>-</sup><br>(pP<sub>*xyI-vipR*</sub>) | Strain HD73<sup>-</sup> expressing *vipR* under the control of the xylose-inducible promoter of *xyIA*. | Km<sup>R</sup> |
| HD73<sup>-</sup><br>(pP<sub>*xyI-vipR*</sub>,<br>pHT-P<sub>*vip3*</sub>/pHT-P<sub>*vipR*</sub>/<br>pHT-P<sub>*vip3-Mut*</sub>) | Strain HD73<sup>-</sup> expressing *vipR* under the control of the xylose-inducible promoter of *xyIA* and used to measure the transcriptional activity of the *vip3A*, *vip3A-Mut* and *vipR* promoter using the *lacZ* reporter gene in the presence of VipR. | Km<sup>R</sup>, Em<sup>R</sup> |
| HD73<sup>-</sup><br>(pHT-P<sub>*vip3-Mut*</sub>) | Strain HD73<sup>-</sup> harboring the pHT-P<sub>*vip3-Mut*</sub> plasmid to measure the transcriptional activity of the *vip3A* promoter when the palindromic sequence is disrupted. | Em<sup>R</sup> |
| HD73<sup>-</sup><br>(pHT-P<sub>*vipR*</sub>) | Strain HD73<sup>-</sup> harboring the pHT-P<sub>*vipR*</sub> plasmid to measure the transcriptional activity of the *vipR* promoter. | Em<sup>R</sup> |
| HD73<sup>-</sup> Δ*spo0A*<br>(pHT-P<sub>*vipR*</sub>) | Strain HD73<sup>-</sup> Δ*spo0A* harboring the pHT-P<sub>*vipR*</sub> plasmid to measure the transcriptional activity of the *vipR* promoter. | Em<sup>R</sup> |
| HD73<sup>-</sup><br>(pHT-P<sub>*vipR-vipR*</sub>) | Strain HD73<sup>-</sup> harboring the pHT-P<sub>*vipR-vipR*</sub> plasmid to measure the transcriptional activity of the P<sub>*vipR*</sub> DNA fragment in the presence of VipR. | Em<sup>R</sup> |
| HD73<sup>-</sup><br>$\Delta spo0A$ (pHT-<br>$P_{vipR-vipR}$) | Strain HD73<sup>-</sup> $\Delta spo0A$ harboring the pHT-$P_{vipR-vipR}$ plasmid to measure the transcriptional activity of the $P_{vipR}$ DNA fragment in the presence of VipR. | Em<sup>R</sup> |
| HD73<sup>-</sup><br>(pHT-$P_{vipR-vipR}$-<br>$P_{vip3}$) | Strain HD73 Cry<sup>-</sup> harboring the pHT-$P_{vipR-vipR-P_{vip3}}$ plasmid to measure the transcriptional activity of the *vip3A* promoter in the presence of a *vipR* expression cassette. | Em<sup>R</sup> |
| HD73<sup>-</sup><br>$\Delta spo0A$ (pHT-<br>$P_{vipR-vipR-P_{vip3}}$) | Strain HD73<sup>-</sup> $\Delta spo0A$ harboring the pHT-$P_{vipR-vipR-P_{vip3}}$ plasmid to measure the transcriptional activity of the *vip3A* promoter in the presence of a *vipR* expression cassette. | Em<sup>R</sup> |
| BL21 (vip3) | *E. coli* BL21(DE3) strain harboring the pET28a$\Omega vip3A$ plasmid, used to produce de Vip3Aa protein. | Km<sup>R</sup> |

### DNA manipulation

Plasmids were extracted from *E. coli* cells by the alkaline lysis method using the Promega DNA Extraction Kit. DNA fragments were purified using Promega Gel and PCR Clean-Up System. Chromosomal DNA was extracted from exponentially growing *B. thuringiensis* HD1 cells using the Qiagen Puregene Yeast/Bacteria kit. Restriction enzymes and T4 DNA ligase were purchased from New England Biolabs and used following the manufacturer’s protocol. The primers used in the study (Table 3) were synthesized by Eurofins Genomics. PCR was performed with a 2720 Thermal cycler (Applied Biosystems) or a Master cycler Nexus X2 (Eppendorf). All the constructs were verified by PCR and sequencing. Sequencing was performed by Eurofins genomics.

**Table 3:**
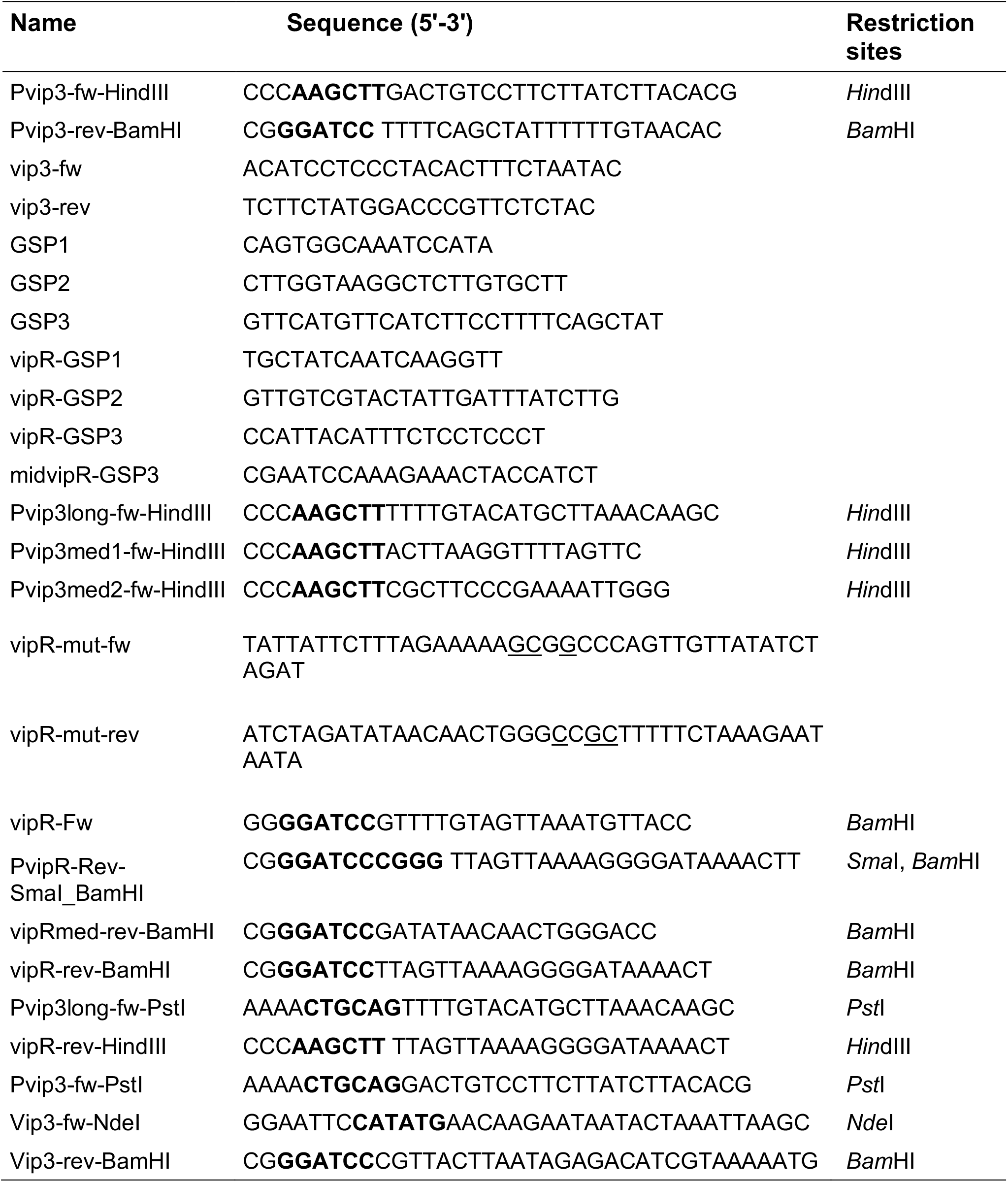
Primers used in the study.

### Bioinformatic analyses

Analysis of the DNA sequence of the HD1 pBMB299 plasmid (NZ_CP004876.1) identified a series of 17 genes, among them the *vip3A* and 3 *cry* genes, all encoded in the same direction, and defined as an island of insecticidal toxins. Each gene of the toxins island of the pBMB299, used as reference, was compared using BLAST (https://blast.ncbi.nlm.nih.gov/Blast.cgi) with the corresponding gene on the *vip3A* plasmid of the eigth following strains: *B. thuringiensis* BGSC4C1 (CP015177), CT-43 (CP001910), HD12 (CP014853), HD29 (CP010091), IS5056 (CP004136), L7601 (CP020005), YBT-1520 (CP004861) and YC-10 (CP011350).

The sequence of the *orf-HTH-encoded* protein (WP_000357137.1) was subjected to structural prediction using the Phyre2 software (27) available online and the HHpred server (54) that detects structural homologues. The intergenic region upstream from the *vip3A* coding sequence was analyzed for the presence of secondary structures in nucleic acid sequences using the Mfold web server (28). The same sequence was also analyzed for the presence of a potential Rho-independent transcription terminator using the ARNold web server (http://rssf.i2bc.paris-saclay.fr/toolbox/arnold/).

### Vip3A protein production and purification

The BL21 (vip3) strain was grown in LB medium supplemented with kanamycin (20 μg/mL) at 37°C to an OD_600nm_ of 0.6. The expression of *vip3A* was induced by adding IPTG (1 mM) and growth was continued for 4 h at 37°C. Bacterial cells were collected by centrifugation and resuspended in 5 % of the initial culture volume in the lysis buffer (50 mM Tris, 300 mM NaCl, 7.5 % glycerol, pH8.0). The bacteria were treated with lysozyme (1 mg/mL) on ice for 60 min and then lysed by sonication. The suspension was centrifuged at 5095 × *g* for 10 min to remove bacterial debris. The supernatant that contains the Vip3A protein was loaded onto 1 mL of Ni-NTA agarose resin previously equilibrated with the lysis buffer. The resin with bound Vip3A proteins was successively washed with 4 mL of lysis buffer containing imidazole (25 and 50 mM). The Vip3A protein was eluted with 1.5 mL of lysis buffer containing imidazole (250 and 500 mM). To remove the imidazole, the buffer of the protein was exchanged against PBS using a PD-10 column (Sigma). The purified protein was stored at −80°C before use.

### Conjugative transfer of the *vip3A*-encoding plasmid pBMB299

The *B. thuringiensis* HD1 (pHT1618K) strain was used as a pBMB299 plasmid donor strain, and the streptomycin resistant HD73^-^ Sm^R^ was used as the recipient strain. The donor and recipient strains were grown in LB at 37°C until OD_600nm_ 0.7. Then 5×10^6^ cells of the donor and recipient strain were mixed in 2 ml BHI broth. The mixed bacteria were transferred on a 0.45 μm membrane by passing the bacterial suspension through the Swinnex® filter holder. The membrane was then put onto a BHI plate and incubated at 37°C overnight. The bacteria were collected by scrapping and resuspended in physiological water. The suspension was then diluted and plated on LB plates containing kanamycin (200 μg/mL) and streptomycin (200 μg/mL) to select the exconjugant bacteria. The presence of the pBMB299 plasmid in colonies that have received the pHT1618K was confirmed by PCR using the primer pair vip3-fw/vip3-rev targeting the *vip3A* gene. The strain HD73^-^ Sm^R^ (pHT1618K, pBMB299) was finally cured of the pHT1618K as described (25). Briefly, the bacterial strain was grown on HCT medium for 3 days at 30°C until sporulation and spores were plated on LB medium containing streptomycin (200 μg/mL). Isolated colonies were screened for the loss of kanamycin resistance on plates. The presence of the pBMB299 in HD73^-^ Sm^R^ Kan^S^ clones was confirmed by PCR using the primer pair vip3-fw/vip3-rev.

### Western blot assays

For western blot, *B. thuringiensis* HD1 and HD73^-^ Sm^R^ (pBMB299) strains were grown in LB medium at 37°C in agitated cultures. For each time point, 50 mL culture was collected. Bacteria were separated from the growth medium by centrifugation at 5095 × *g* for 10 min. The bacterial cell pellet was resuspended in the lysis buffer (50 mM Tris, 300 mM NaCl, 7.5% glycerol, pH8.0). The suspension was then treated with lysozyme (1 mg/mL) on ice for 1 h, and sonicated. The proteins of the culture medium were precipitated according to the following steps: dithiothreitol (100 mM) was added to the suspension to prevent protein oxidation, then (NH4)2SO4 (19.62 g) was slowly added to 45 mL of sample to reach 70% saturation (0°C) and the suspension was incubated overnight with slow agitation at 4°C. The precipitated proteins were collected by centrifugation at 5095 × *g* for 10 min, and resuspended in 200 μL lysis buffer. The protein concentration of the pellets and the precipitated supernatant samples was determined using the Bradford method (55). 20 μg proteins of each sample and 0.05 μg purified Vip3A were separated using SDS-PAGE on a 7.5% polyacrylamide gel. Gels were either stained with Coomassie brilliant blue or subjected to Western blot assays. For Western blot assays, the proteins were electro-transferred to a PVDF membrane (Immun-Blot® PVDF Membrane, Bio-Rad). The membrane was blocked for 1 h in 5% skim milk dissolved in TBS-T buffer (10 mM Tris-HCl, 150 mM NaCl, 0.05% Tween-20, pH 8.0), then treated with anti-Vip3Aa11 polyclonal antibodies (56) diluted 1:100 000 in 5% skim milk TBS-T buffer for 1 h at room temperature. The membrane was washed three times with 15 mL TBS-T buffer, and then treated with 1:20 000 diluted Goat-anti-rabbit antibodies (Invitrogen G21234) for 1 h. Membranes were washed three times with TBS-T buffer before being revealed using the SuperSignal^TM^ West Pico Chemiluminescent substrate (ThermoScientific) according to the manufacturer’s instructions, and imaged using the Chemidoc system (Bio-Rad).

### ß-galactosidase assay

The ß-galactosidase activity was monitored using a qualitative and quantitative method. For the qualitative assay, cells were streaked on LB plates containing X-gal (50 μg/mL) and incubated at 37°C for the indicated periods of time. The blue coloration reflects the activity of the ß-galactosidase. For the quantitative method, strains were precultured in 10 mL LB medium until OD_600nm_ 0.6-1.0 from freshly isolated strains on plates. Then, the preculture was used to inoculate 50 mL LB medium at an OD_600nm_ = 0.005 and bacteria were allowed to grow until being harvested by centrifugation at the indicated time points. The activity was measured as previously described and expressed as units per milligram of protein (57).

### mRNA extraction and determination of the *vip3a* and *vipR* mRNA 5’-ends

RNA was extracted from the *B. thuringiensis* HD1 strain grown at 37°C in LB medium and harvested at T2. RNA samples were prepared as described (7) and stored at −80°C before use. RACE-PCR was performed using the 5′ RACE System (Invitrogen, cat: 18374058) according to the instruction manual. The cDNA was generated using the specific primer GSP1. The subsequent PCR steps were realized with the nested gene-specific GSP2 and GSP3 primers. The 5’-end of *vip3A* mRNA was determined by sequencing the PCR product using GSP3. For *vipR* mRNA 5’-end determination, RNA was extracted from the *B. thuringiensis* HD73^-^ strain grown at 37°C in LB medium and harvested at T2. The cDNA was generated using the specific primer vipR-GSP1. The subsequent PCR steps were realized with the nested gene-specific vipR-GSP2 and vipR-GSP3 primers. The *vipR* mRNA 5’-end P1 was determined by sequencing the PCR product using the primer vipR-GSP3 and P2 was identified by sequencing the PCR product using the primer midvipR-GSP3.

## Acknowledgments

We are grateful to the Prof. Jie Zang and Dr Zeyu Wang for the kind gift of the anti-Vip3Aa antibody. We thank Dr Michel Gohar for his help with the long-term cultures. We thank Alicia Nevers for the Northern blot experiments.

## Data availability statement

Data are available on request from the authors.

## Notes

### Competing Interest Statement

The authors have declared no competing interest.

