## Supplemental Fig. S1, S2, S3, S4, and Table S1 for "Expression of the *Bacillus thuringiensis vip3A* insecticidal toxin gene is activated at the onset of stationary phase by VipR, an autoregulated transcription factor"

Running title: Regulation of *vip3A* expression

*Present address: School of Basic Medical Science, Xiangnan University, Chenzhou, Hunan, 423000, PR China

Key-words: Bacillus, biopesticide, gene regulation, plasmid, transcription

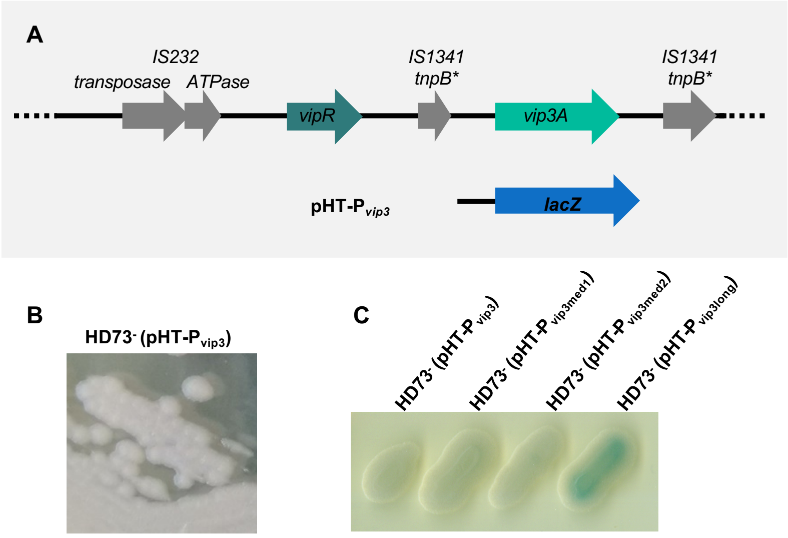


**Fig. S1- Vip3A is not expressed in the HD73$^-$ strain.**

**A-** Genetic organization of Bt HD1 pBMB299 plasmid region containing the *vip3A* gene (NZ_CP004876.1). The asterisk indicates a gene containing nonsense mutations. Schematics representing the DNA fragment screened for its ability to produce a transcriptional activity. **B-** The HD73$^-$ (pHT-P$_{vip3}$) strain was isolated on LB X-gal (50 µg/mL) plates and grown at 37°C for 24 h. **C-** The HD73$^-$ strains carrying the pHT-P$_{vip3}$, the pHT-P$_{vip3\ med1}$, the pHT-P$_{vip3med2}$, or the pHT-P$_{vip3long}$ were isolated on LB X-gal (50 µg/mL) plates and grown at 37°C for 24 h.

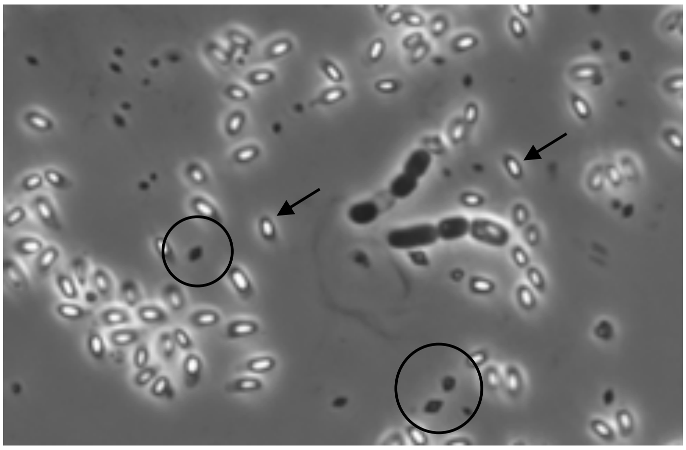

**Fig. S2- Conjugative transfer of the pBMB299 into the HD73$^{-}$ strain allows the cells to produce Cry crystals.** Phase contrast microscope images of Bt HD73$^{-}$ Sm$^{R}$ (pBMB299) sporulating cells. Cells were grown on HCT plates for 4 days at 30°C. Spores are the refringent structures indicated with arrows. The crystals are indicated using circles.

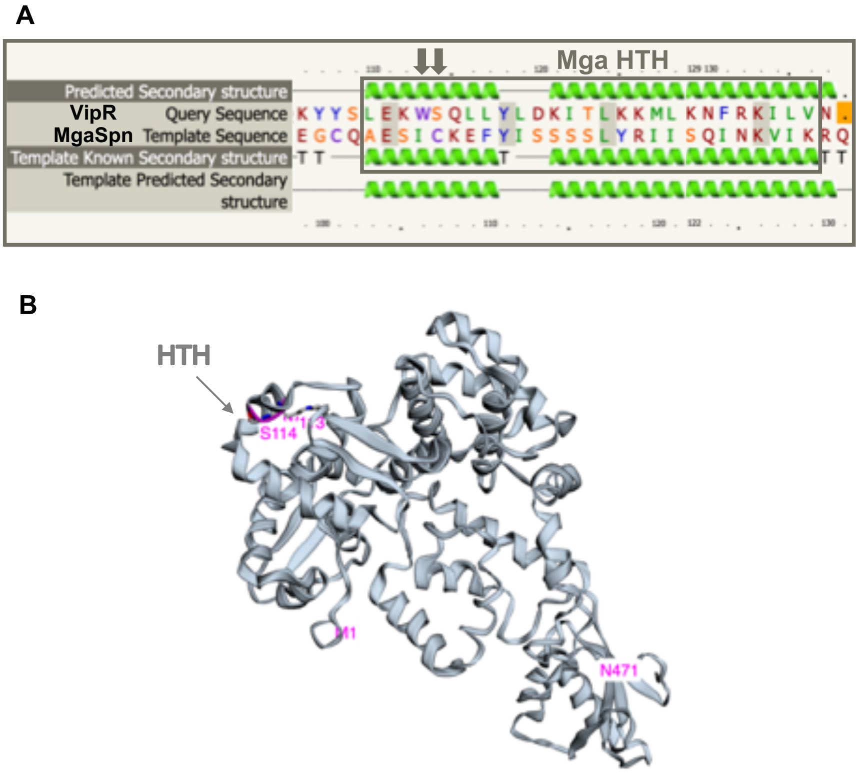


**Fig. S3- Mutation of the VipR HTH. A-** Alignment of the VipR and MgaSpn protein sequences. The sequence corresponding to the HTH domain of *S. pyogenes* Mga of is boxed. Arrows point to the 2 amino acids that were selected to be mutated. **B-** 3D model of the VipR structure modelized by the Phyre2 webserver. The W113 and S114 AA are indicated in pink. The M1 and N471 amino acids at the N- and C-termini are indicated.

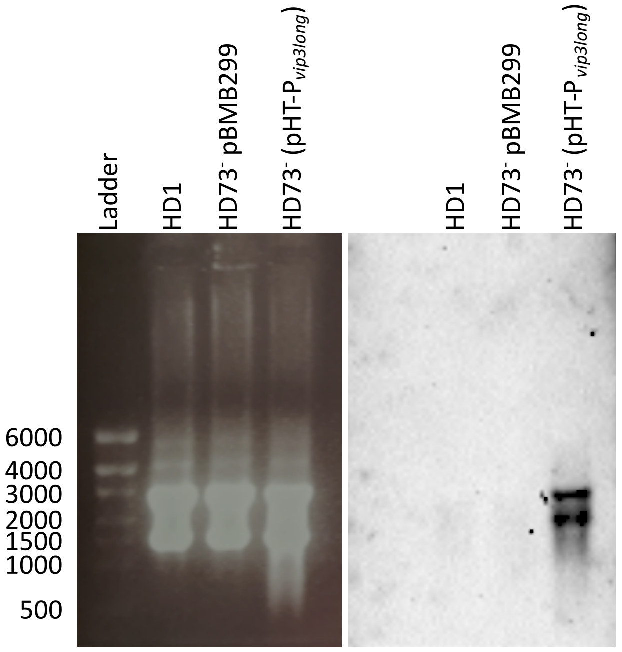


**Fig. S4 Northern Blot analysis of *vipR* trancscription indicates the presence of two transcripts in the HD73$^{-}$ (pHT-P$_{vip3long}$) strain.** 10 µg RNAs were separated on a 1% agarose gel and transferred overnight by capillarity on a nylon membrane (GE Healthcare, #RPN303B) in SSC 10X Buffer. RNAs were then UV-crosslinked to the membrane using a Stratalinker apparatus. Generation and incubation of the membrane with DNA dig-labeled probes was performed using the Dig-High Prime DNA Labeling and Detection Starter Kit II (Roche, #11585614910), following manufacturer's instructions. Primers used to generate the DNA probes are NB-vipR-fw (GCATAAGTTCAATTATATGCGAATTG) and NB-vipR-fw (TTTCAGAAGATATTGTTTGGAATAAATGT). Membranes were washed twice in 2X SSC 0.1% SDS and once in 1X SSC 0.1% SDS, for 10 min at 65 °C. Revelation was done according to the kit instructions using a Chemidoc System (Bio-Rad).
Numbers indicated at the left indicate the size (in bases) of the single-stranded RNA transcripts used in the RiboRuler High Range RNA Ladder (Thermo Scientific).

### Table S1- Comparison of the genes in the *vip3A* locus from nine Bt strains

| Bt strains | bp | HD1 | YBT-1520 | CT-43 | L-7601 | IS5056 | HD-29 | BGSC 4C1 | HD-12 | YC-10 |
|---|---|---|---|---|---|---|---|---|---|---|
| Plasmid Accession No. | | NZ_CP004876 | CP004861 | CP001910 | CP020005 | CP004136 | CP010091 | CP015177 | CP014853 | CP011350 |
| IS232 family transposase | 1296 | 100 | 100 | 100 | | 100 | 100 | | | 100 |
| IS232 helper ATPase | 753 | 100 | 100 | 100 | | 100 | 100 | | | 100 |
| HTH-containing protein | 1416 | 100 | 100 | 100 | 95 (1343/1416) | 100 | 99 (1413/1416) | 99 (1408/1416) | 90 (1270/1416) | 100 |
| transposase | 532 | 100 | 99 (529/532) | 99 (529/532) | 91 (191/210) | 99 (529/532) | 99 (529/532) | 98 (524/532) | 91 (191/210) | 99 (539/532) |
| vip3A | 2370 | 100 | 100 | 100 | 89 (1791/2013) | 100 | 100 | 99 (2358/2370) | 96 (1967/2053) | 100 |
| transposase | 929 | 100 | 100 | 100 | 85 (809/948) | 100 | 100 | 99 (917/930) | 86 (822/953) | 100 |

The Bt HD1 gene sequences were used as the reference for the comparison with the gene of the other Bt strains.
The numbers in the table represent the percentage of identity of each gene obtained by Blast comparison. Numbers between parentheses indicate the number of base pairs that are identical to the corresponding Bt HD1 gene. Empty case means that no homolog was found.
